# Rational Design Reveals Structural Plasticity of the CsgA β-Solenoid Enabling Programmable Autogenic Engineered Living Materials

**DOI:** 10.64898/2026.02.23.707537

**Authors:** Hoda M. Hammad, Seth Swarnadeep, Harrison Priode, Erin C. Jackson, Anna Kurowski, Robert B. Moore, Avinash Manjula-Basavanna, Sanket A. Deshmukh, Anna M. Duraj-Thatte

## Abstract

Microbial functional amyloids play central roles in biofilm formation and serve as foundational building blocks for autogenic engineered living materials (ELM), yet the structural design space governing their assembly and stability remains poorly defined. In *Escherichia coli*, the β-solenoid protein CsgA functions as a canonical extracellular matrix scaffold, but prior engineering efforts have primarily focused on terminal functionalization rather than modification of the β-solenoid core itself. Here, inspired by the evolutionary diversification of CsgA-like proteins, which expands along the vertical fiber axis, we explore a second orthogonal axis of structural plasticity: the horizontal dimension of the β-solenoid. We rationally designed a library of CsgA variants in which the length of individual β-strands was systematically reduced or extended from the native seven residues to lengths spanning 3–21 residues, while preserving conserved gate residues and loop regions. Integration of AI-based structure prediction using AlphaFold2 with all-atom molecular dynamics simulations reveals that β-solenoid stability arises from a balance among strand length, residue composition, and solvent interactions, thereby defining both lower and optimal bounds for nanofiber assembly. Experimental validation demonstrates that engineered *Escherichia coli* can secrete and assemble these CsgA variants into extracellular nanofibers through the native curli biogenesis machinery, while preserving the characteristic cross-β architecture. Additionally, the CsgA β-solenoid variant library translates molecular design into macroscopic ELM, with deletion variants showing an inverse relationship between stiffness and extensibility, from highly extensible 3aa to stiff, strong 5aa films. Insertion-based variants largely retain CsgA-like extensibility while enabling tunable stiffness and strength across strand lengths. Together, these findings uncover previously unrecognized structural plasticity in microbial β-solenoid proteins and establish β-strand length as a generalizable design parameter linking molecular architecture to nanofiber stability, with implications spanning microbial functional amyloids and the rational design of autogenic engineered living materials.

## Introduction

Engineered living materials (ELMs) are an emerging class of materials that integrate living cells to generate dynamic and functional materials with applications in sustainability, biomedicine, biosensing, and environmental remediation. (1) Among these, autogenic ELMs leverage engineered cells to produce and assemble a functional polymeric matrix in situ. (2) This approach enables capabilities that are difficult to achieve with conventional biomaterials and exogenic ELMs (in which the polymeric matrix is sourced externally), including self-assembly, self-regeneration, environmental responsiveness, and on-demand functionalization. (3, 4) Despite these advantages, autogenic ELMs remain comparatively underexplored, with existing systems constrained primarily by a narrow range of genetic engineering strategies.(5)

Among the key components of autogenic ELM are extracellular matrix (ECM) protein nanofibers produced by *Escherichia coli* that self-assemble from the β-solenoid protein CsgA, which have been widely studied.(6, 7) These protein nanofibers, originating from microbial biofilms, exhibit exceptional thermal and chemical stability, high mechanical strength, and tunable functionality enabled through genetic engineering.(2, 8) Over the past decade, engineering efforts have predominantly focused on the C-terminal fusion of functional peptides or proteins to CsgA to achieve specific applications, such as biosensing, catalysis, or biomolecular binding.(4, 9–11) To a lesser extent, N-terminal and dual-terminal fusion strategies have been explored to enable spatially controlled functional display and to support multifunctional scaffolds.(12–15) In contrast, direct engineering of the CsgA β-solenoid core itself, which governs nanofiber self-assembly, stability, and intermolecular packing, has remained largely unexplored, with only a small number of studies reporting internal repeat deletions or residue-level modifications within the CsgA β-solenoid domain.(13, 16–18)

In our previous work, we examined the evolutionary diversification of CsgA-like β-solenoid proteins across bacterial species and identified a consistent trend in which β-solenoid length expands vertically along the fiber axis.(2) This diversification increases the number of β-strand repeat units within CsgA-like β-solenoid proteins, enabling the formation of longer CsgA monomers with enhanced structural and functional capabilities. Motivated by this observation, we pursued a complementary design rationale: rather than extending the protein along the vertical axis, as observed in nature, we asked whether the horizontal axis of the β-solenoid, specifically the length of individual β-strands, could be rationally engineered. Modulating β-strand length represents an orthogonal strategy for tuning β-solenoid architecture and may reveal new structure-function relationships relevant to autogenic ELMs.

The β-solenoid structure of CsgA consists of repeating β-strands that stack head-to-tail to form curli nanofibers, an architecture that is fundamental to both nanofiber stability and the functionalization of nanofiber surfaces.(19, 20) To date, the β-solenoid core itself has remained largely unexplored as a design variable, despite evidence of substantial evolutionary flexibility in β-solenoid structures. Directly engineering the structural β-solenoid core offers an opportunity to define microbial β-solenoid structural plasticity and clarify how molecular-level variations in β-solenoid design translate into emergent physicochemical properties of ELMs. More broadly, it could provide a future roadmap for rationally tuning the physical, chemical, and biological properties of ECM-based materials.

In this work, we expand the scope of CsgA engineering by designing variants that modulate the horizontal dimension of the CsgA monomer, thereby directly altering the core β-solenoid architecture. Specifically, we increased the number of amino acids per β-strand, extending the β-strand length from the native seven residues per β-strand to variants with lengths ranging from 3 to 21 residues. By systematically varying the hydrophobic and hydrophilic composition within the β-strands while preserving conserved terminal regions, we investigated how modifications to the β-solenoid core affect nanofiber assembly and stability, thereby providing new insights into the design principles governing β-solenoid proteins. To support this approach, we employed the artificial intelligence-based structure prediction tool AlphaFold2 (21) to model β-solenoid variants and evaluated the stability of the resulting folds using all-atom (AA) molecular dynamics (MD) simulations in explicit solvent. Leveraging the curli secretion machinery of *E. coli,* as previously reported (2), we show that these engineered CsgA variants retain the ability to self-assemble into curli nanofibers and macroscopic ELM (**Figure 1**).

**Figure 1.**
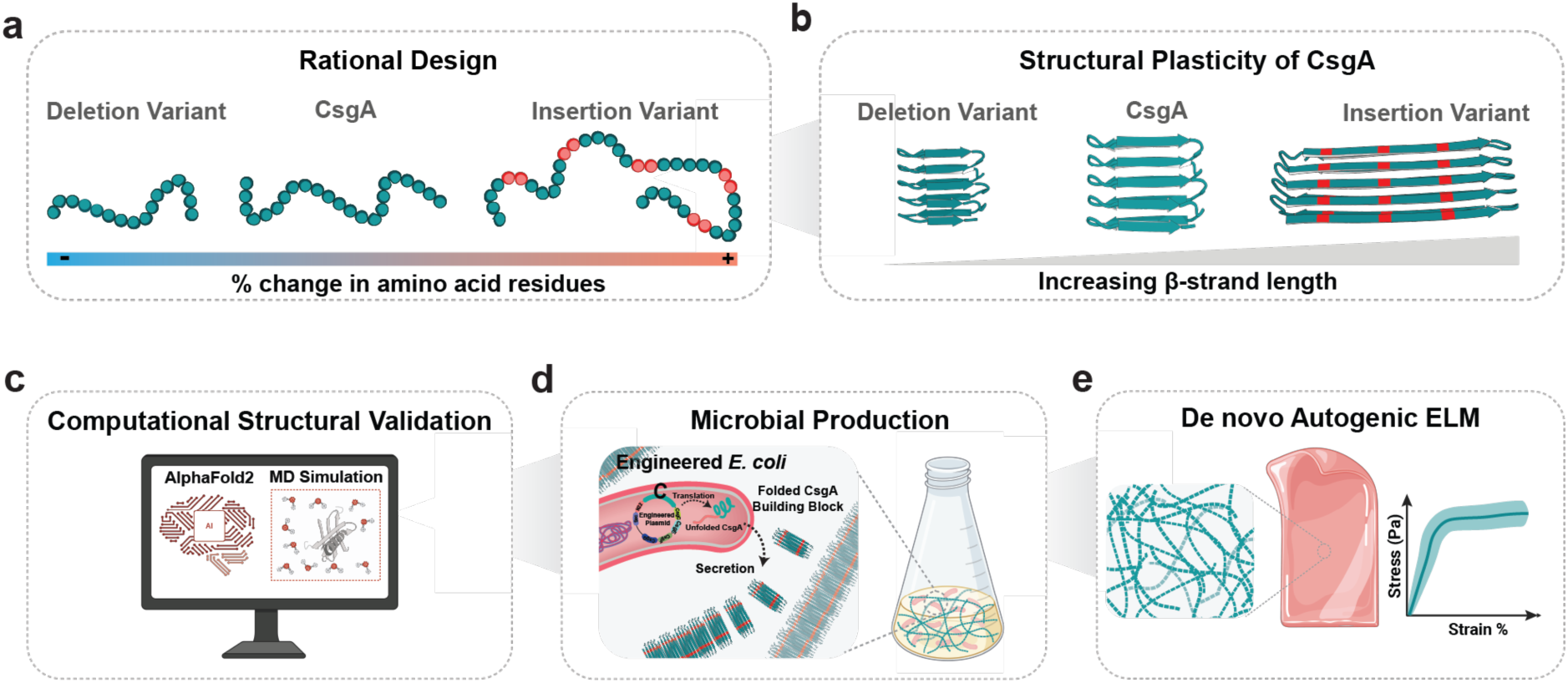
Schematic summary of rationally designed β-Solenoid Protein CsgA for de novo Autogenic ELM.

## Results and Discussion

### Rational Design of the Horizontal Dimension of CsgA β-Solenoid Protein

Previous studies have identified an evolutionary trend in which CsgA-like β-solenoid proteins from diverse organisms expanded along the vertical axis by increasing the number of β-strand repeats, reaching up to 46 repeats, compared with the 5 repeat units present in the wild-type *E. coli* CsgA. (2)These naturally occurring variants exhibit distinct macroscopic physicochemical properties, indicating that changes in β-solenoid architecture directly influence the mechanical and functional properties of the resulting ECM. Building on this observation, we investigated whether the horizontal axis of the CsgA β-solenoid core, specifically the length of individual β-strands, could be systematically engineered to modulate nanofiber assembly.

The β-solenoid core of *E. coli* CsgA is composed of five repeating antiparallel β-strand units, each defined by a conserved 7-residue motif: KR_1_–Ω_L_–Ψ_L_–Ω_C_–Ψ_R_–Ω_R_–KR_7_. The terminal key residues (KR_1_ and KR_7_) play a critical role in stabilizing strand orientation and promoting head-to-tail stacking during nanofiber assembly.(2, 22) The inward-facing Ψ residues, which are enriched in hydrophobic or weakly polar amino acids (e.g., A, I, V, L, F, S, T), form the interior core of the β-solenoid, whereas the outward-facing Ω residues are generally more hydrophilic and solvent-exposed, defining the exterior surface of the fibril. Adjacent antiparallel β-strands are connected by conserved intra- and inter-repeat loop regions, most characterized by an XGXX consensus motif (**Figure 2a**). (2, 22, 23)

**Figure 2.**
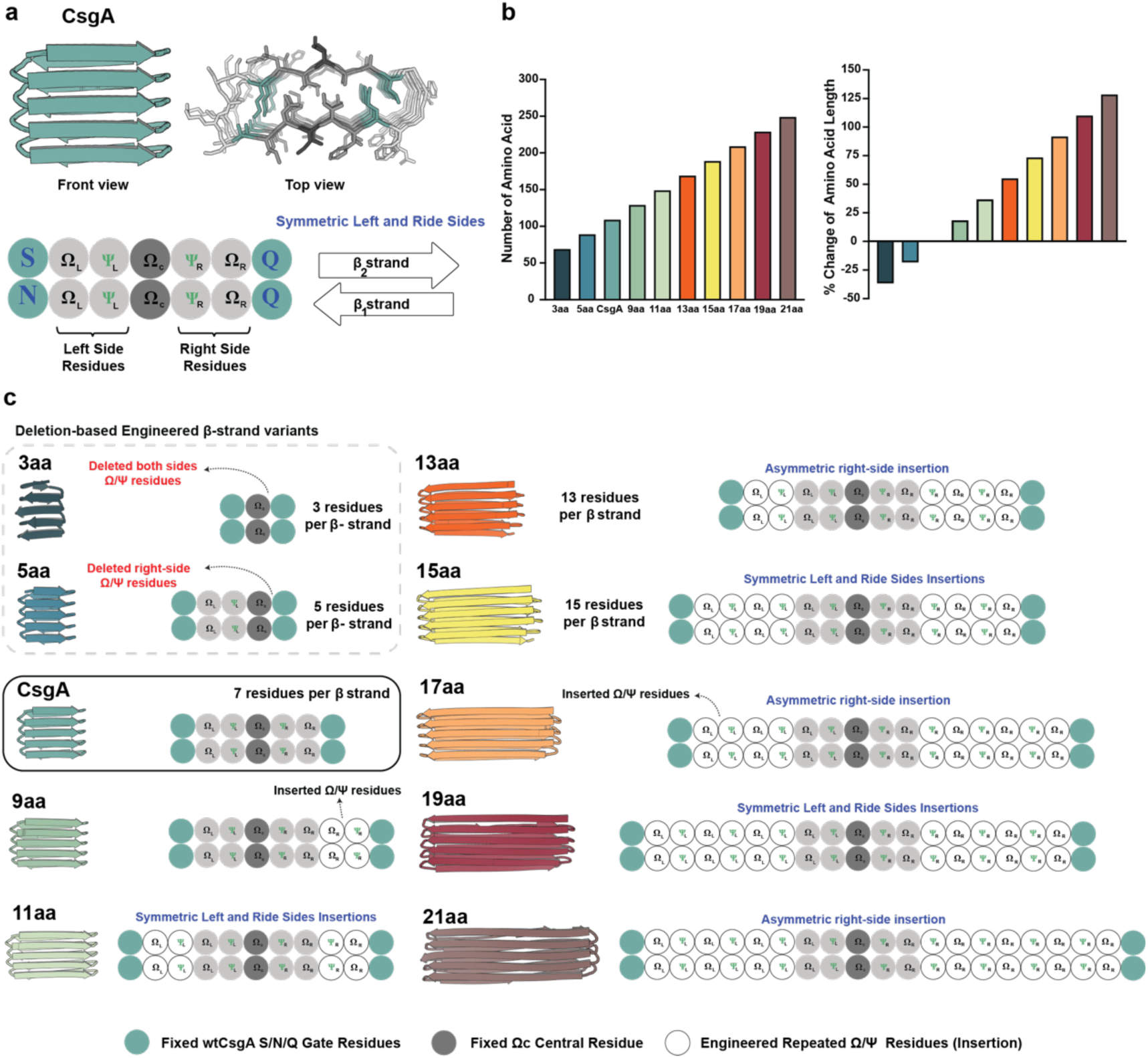
Structural Plasticity of the CsgA β-Solenoid via Rational β-Strand Length Design. **a.** Wild-type CsgA β-Solenoid architecture, **b.** Number of amino acids in the designed β-Solenoid variants, **c.** Library of rationally designed β-Strand Length variants.

Using this canonical organization as a reference, we rationally designed a library of CsgA β-strand variants that systematically vary β-strand length while preserving the native β-solenoid framework. In all designs, the gate residues (KR_1_ and KR_7_), the central outward-facing residue (Ω_C_), and all intra- and inter-repeat loop sequences were left unchanged. Variants were generated by modifying the left (Ω_L_–Ψ_L_) and/or right (Ψ_R_–Ω_R_) Ω/Ψ positions flanking Ω_C_, thereby altering β-stand length without disrupting conserved β-solenoid structural features.

Across the design space, we generated two classes of variants relative to the wild-type 7-residue β-strand, CsgA: 1) deletion-based variants and 2) insertion-based variants. The resulting library comprises 3aa, 5aa-L, 5aa-R, 9aa-L, 9aa-R, 11aa, 13aa-L, 13aa-R, 15aa, 17aa-L, 17aa-R, 19aa, and 21aa-L and 21aa-R, in addition to CsgA (**Figure 2b**). Those odd-number variants ensure symmetric presentation of hydrophobic and polar residues across the two faces of the β-solenoid structure. Here, “aa” denotes the total number of amino acids per β-strand, while the “L” or “R” suffix indicates whether Ω/Ψ pairs were modified on the left or right side of Ω_C_, respectively.

### Deletion-based designs

Deletion-based variants were designed by selectively removing Ω/Ψ pairs (Ω_L_–Ψ_L_ and/or Ψ_R_–Ω_R_) from each β-strand of CsgA. For the 3aa variant, both the left and right Ω/Ψ positions (Ω_L_–Ψ_L_ and Ψ_R_–Ω_R_) were deleted, leaving the minimal β-strand motif KR_1_–Ω_C_–KR_7_, which preserves the key residues and the central outward-facing residue (Ω_C_). For the 5aa variants, two deletion patterns were designed: a left-deleted variant (5aa-L), lacking Ω_L_–Ψ_L_, and a right-deleted variant (5aa-R), lacking Ψ_R_–Ω_R_. In all cases, the conserved key residues KR_1_, KR_7_, and Ω_C_ were retained.

### Insertion-based designs

Insertion-based variants were generated by repeating Ω/Ψ pairs on the left and/or right sides of the conserved Ω_C_ position. Each insertion corresponded to the addition of one Ω–Ψ pair, repeated either on the left side (Ω_L_–Ψ_L_) or the right side (Ψ_R_–Ω_R_) of each β-strand. For the 9aa variants, a single Ω/Ψ pair was added asymmetrically, producing 9aa-L (left repetition) and 9aa-R (right repetition). In the 11aa variant, Ω/Ψ pairs were added symmetrically, resulting in two Ω/Ψ pairs on each side of Ω_C_, yielding a single variant (11aa). The 13aa variants were generated by asymmetrically adding one additional Ω/Ψ pair to the 11aa design, yielding 13aa-L and 13aa-R. For the 15aa variant, symmetric repetition was applied to the 13aa variant, resulting in four Ω/Ψ pairs on each side of Ω_C_. The 17aa variants were generated by asymmetrically adding one Ω/Ψ pair to the 15aa variant, producing 17aa-L and 17aa-R. For the 19aa variant, symmetric repetition was again applied, producing six Ω/Ψ pairs on each side of Ω_C_. Finally, the 21aa variants were designed by asymmetrically adding an additional Ω/Ψ pair to the 19aa strand, yielding 21aa-L and 21aa-R.

In total, this rational design strategy generated three deletion-based variants and eleven insertion-based variants by systematically redistributing Ω/Ψ pairs around a conserved central Ω_C_ while maintaining the native β-solenoid framework. These designs alter the number and spatial arrangement of inward- and outward-facing Ω/Ψ residues without modifying gate residues or loop regions. All variants were subsequently evaluated using AlphaFold2 for three-dimensional structure prediction to assess retention of the β-solenoid fold (**SI Appendix Fig. S1 and S2**). For strand lengths with both left and right variants, the model with higher-confidence prediction was selected for further analysis (SI Appendix Fig. S1), resulting in a final panel of ten variants, CsgA plus nine engineered variants (3aa, 5aa-R, 9aa-R, 11aa, 13aa-R, 15aa, 17aa-R, 19aa, and 21aa-R). These variants were carried forward for computational and experimental characterization and are hereafter referred to solely by β-strand length to simplify nomenclature.

### Physicochemical Properties of β-Solenoid Protein Variants

Additional analyses were performed to characterize the physicochemical properties of CsgA and the nine engineered β-strand variants (Supplementary Table S1 and Table S2). As expected, the total number of amino acid residues increased monotonically with β-strand length, ranging from 131 residues for CsgA to 271 residues for the 21aa variant, with a corresponding increase in molecular weight from 13.1 kDa to 27.2 kDa. Increasing β-strand length led to a progressive increase in net negative charge, shifting from −2e for the shortest variant (3aa) to −19e for the longest variant (21aa), compared to −6e for CsgA. Across all variants, the fraction of positively charged residues remained relatively constant (∼2.7–3.3%), whereas the fraction of negatively charged residues increased modestly with strand length, resulting in a gradual increase in total charged residue content from 8.8% to 12.9% (**SI Appendix Table S1**). In addition, the predicted hydrophobicity, as quantified by the GRAVY score (**SI Appendix Fig. S3, Table S1**), varied systematically with β-strand length. Shorter variants exhibited more negative GRAVY values (e.g., 3aa and 5aa), indicating increased overall hydrophilicity. In contrast, longer variants showed progressively higher GRAVY scores approaching neutrality or slight hydrophobicity, consistent with the cumulative addition of Ω/Ψ residue pairs within the β-solenoid core. These trends are consistent with the underlying design strategy, in which β-strand length was increased by repeating Ω/Ψ residue pairs rather than by introducing new charged motifs. Because Ψ positions are typically occupied by hydrophobic or weakly polar residues, whereas Ω positions are more frequently solvent-exposed and enriched in polar or charged residues, increasing β-strand length redistributes outward- and inward-facing (Ω/Ψ) residues while largely preserving the intrinsic charge balance of the β-solenoid scaffold.

### Molecular Basis of Structural Stability in β-Solenoid Protein Variants

To evaluate the structural stability of the β-Solenoid protein variants in aqueous solution, we computed the root-mean-square deviation (RMSD) of the C_α_ atoms of β-strand residues relative to their AlphaFold2-predicted starting conformations. RMSD profiles for CsgA (7aa) and nine engineered variant systems are shown in **Figure 3a**, with the legend showing the average RMSD over the final 250 ns of a 500 ns molecular dynamics (MD) simulation. CsgA exhibits a moderate, relatively stable RMSD of 5.9 ± 1.6 Å, serving as a baseline for comparison. Several variants, including 5aa, 9aa, 13aa, and 21aa, exhibit slightly lower RMSDs, while 11aa, 15aa, and 19aa show values comparable to CsgA, suggesting that these systems retain similar β-strand architecture during the simulations. Notably, 5aa demonstrates the lowest RMSD (1.3 ± 0.3 Å), indicating an exceptionally stable and rigid conformation. In contrast, the 3aa and 17aa variants display significantly elevated RMSDs (9.6 ± 2.1 Å and 9.1 ± 0.6 Å, respectively), indicating pronounced deviations from their initial structures. For 3aa, this likely reflects unfolding, whereas for 17aa, it may result from intermittent flipping of the C-terminal R5 strand (**SI Appendix Fig. S4 and S5**). Overall, all the variants except 3aa were able to maintain stable β-strand percentage throughout the 500 ns simulation (**SI Appendix Fig. S6**).

**Figure 3.**
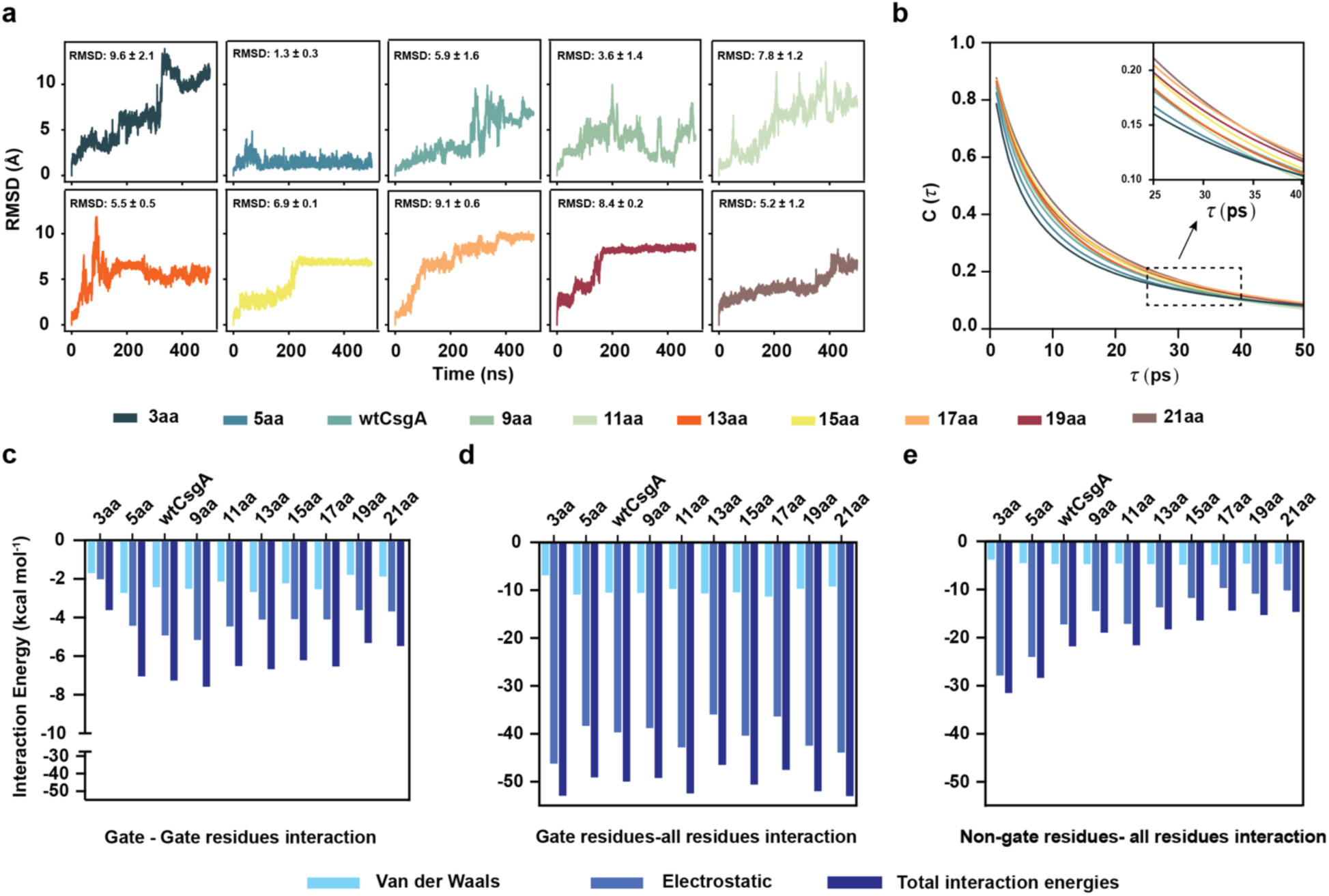
Structural Stability in β-Solenoid Protein Variants. **a.** RMSD of the β-sheets C-alpha atoms of the CsgA variants in water. The average RMSD is calculated from the last 250 ns of simulation trajectories. **b.** Hydrogen bond autocorrelation function among the protein residues. The bar plots show the VdW, electrostatic, and total energies measured between **c**. gate-to-gate residues, **d.** gate-to-all residues, and **e.** non-gate-to-all residues.

Additionally, to evaluate the stability of β-strand architecture and backbone conformational preferences in the CsgA and nine variants, Ramachandran plots of backbone dihedral angles (*ϕ* and *ψ*) were generated for all residues (**SI Appendix Fig. S7b**). In CsgA and most variants, excluding 3aa, the plots exhibit a dominant density cluster centered around the reported β-strand region (*ϕ* ≈ −120°, *ψ* ≈ 120°), indicating persistent β-sheet geometry throughout the simulations. The 3aa variant exhibits a broadened distribution, with substantial populations outside the β-strand region, reflecting pronounced backbone disorder and secondary-structure loss. Notably, additional density near *ϕ* ≈ −60° and *ψ* ≈ 125°, characteristic of type II β-turns(24), was observed in some systems, highlighting local structural motifs that mediate inter-strand connectivity. Overall, the Ramachandran analysis supports the conclusion that most variants preserve β-sheet integrity, while 3aa undergoes significant conformational destabilization.

To investigate the flexibility of the variants, the persistence length of individual β-strands was calculated. We found that β1 (N-terminal) and β10 (C-terminal) exhibited lower persistence lengths compared to the core strands (**SI Appendix Fig. S8**). Furthermore, the persistence length increases with β-strand length, consistent with the conformational fluctuations observed in **Figure 3a**. Additionally, residue-wise flexibility in aqueous environments was assessed by calculating the root-mean-square fluctuation (RMSF), as shown in **SI Appendix Fig. S7a**. The analysis reveals that most systems exhibit relatively low fluctuations in their β-strand core regions, indicating preserved structural rigidity—except for 3aa, which stands out with markedly elevated flexibility. As expected, terminal residues across all systems show increased RMSF values due to solvent exposure and the absence of a stabilizing secondary structure. At the N-terminus, the first ∼22 residues adopt a flexible coil-like conformation, contributing to heightened mobility. Additionally, occasional conformational changes in the R5 β-strand contribute to increased fluctuations at the C-terminal end. The 3aa variant, in particular, displays elevated flexibility throughout the entire sequence, consistent with its high RMSD and suggestive of significant unfolding.

To test whether hydrogen bonds within the β-sheets of variants with varying strand lengths further contribute to overall protein structural stability, the hydrogen bond autocorrelation function, C(τ), which captures the temporal persistence of hydrogen bonds between residues within the protein variant, was calculated (**Figure 3b**). A slower decay in C(τ) indicates more stable, longer-lived hydrogen bonds, whereas a faster decay suggests more transient and unstable interactions.(25–27) Among all systems, the 3aa variant exhibits the fastest decay, reflecting highly dynamic and short-lived hydrogen bonds, consistent with its structurally unstable and partially unfolded conformation observed in previous analyses. In contrast, the other variants generally exhibit slower decay with increasing residue numbers, reflecting stronger and more persistent hydrogen-bonding networks that likely contribute to their enhanced structural stability.

### Influence of Key Residues and Electrostatic Forces on β-Solenoid Variants Architecture

To explore the role of key residues, non-bonded interaction energies (IE) between key residues (KR) and all residues were calculated using our custom-developed code. Key residues (KR), Asn and Gln, or Ser, positioned at both termini of each β-strand, were identified as a conserved feature across all variants, with their numbers consistent in each case. Non-key residues (non-KR) were defined as all other residues not located at these key residue positions. Panels in **Figure 3 c-e** present the van der Waals (vdW), electrostatic, and total interaction energies for between KR and KR (IE1), between KR and all residues (IE2), and between non-KR and all residues (IE3) interactions across the CsgA and its variants, with a detailed breakdown provided in **SI Appendix Table 4**. The electrostatic interactions in IE1 consistently exhibit favorable (i.e., negative) interaction energies, underscoring their role in structural stabilization. Among all systems, the 3aa variant shows the weakest KR − KR stabilization, consistent with its overall structural instability, whereas the other variants display total IE1 interaction energies comparable to the CsgA (−7 kcal mol^-1^). The electrostatic interactions in IE2 are significantly stronger (−50 kcal mol^-1^) than IE1 interactions, highlighting the central role of key residues in stabilizing the protein core. In 3aa, the strength of these interactions appears to stem from structural unfolding rather than organized stabilization. In particular, total IE1 interaction energy for 3aa decreased from –6 kcal.mol^-1^ for the first 50 ns (**SI Appendix Fig. S9**) to –4 kcal, mol^-1^ for the last 50 ns while IE2 interaction energy remained unaltered. The electrostatic interactions in IE3, although slightly weaker, remain favorable across all variants. However, their stabilizing effect diminishes with increasing length, likely due to the accumulation and spatial clustering of charged residues, which can introduce electrostatic repulsion and disrupt local packing (**SI Appendix Fig. S10**). Notably, electrostatic interactions dominate the total interaction of energy in all cases, particularly among key residues. Collectively, these results emphasize the importance of electrostatic stabilization, especially involving key residues in preserving β-solenoid structural integrity, and suggest that disruption of these interactions, as in the 3aa variant, may contribute to enhanced conformational disorder.

To investigate the influence of charged residues and electrostatic interactions on β-solenoid protein stability, we performed two sets of additional simulations in which partial charges were neutralized for (a) only charged residues and (b) all residues. The RMSD analysis **(SI Appendix Fig. S11)** revealed that neutralizing the partial charges of all residues significantly disrupted the core β-solenoid secondary structure across all variants. In contrast, neutralizing only the partial charges of charged residues did not destabilize the structure for most variants, except for 5aa, which showed slight destabilization. Notably, enhanced stability was observed for variants 15aa to 21aa, suggesting that neutralizing charged residues mitigates unfavorable electrostatic interactions arising from like-charged residues.

### Mapping the Surface Hydration Landscape of Engineered β-Solenoid Variants

The extent of hydration was assessed by analyzing the radial distribution function (RDF) between the backbone heavy atoms of each variant and the oxygen atoms of surrounding water molecules, as shown in **Figure 4a**. The positions of the first, second, and third RDF peaks are conserved mainly across all variants, including CsgA, indicating that the spatial organization of water around the backbone remains similar in terms of radial distance. However, a notable difference arises in peak intensities, which reflect the degree of local hydration and ordering of water molecules. Specifically, when comparing RDFs from the initial 50 ns of the simulation (**SI Appendix Fig. S12**) to the final 50 ns (**Figure 4a**), the 3aa variant exhibits a substantial increase in the heights of both the first and second peaks. This heightened hydration suggests enhanced water structuring near the protein backbone, consistent with partial unfolding of the β-solenoid architecture. This unfolding of 3aa, which begins around 200 ns (see **Supplementary Movie**), is likely driven by the destabilization of interactions within itself. The resulting disruption of the hydrophobic core permits increased water infiltration, thereby increasing the local hydration density. In contrast, other variants maintain relatively stable peak heights over time, indicating persistent structural integrity and limited water access to the β-solenoid core.

**Figure 4.**
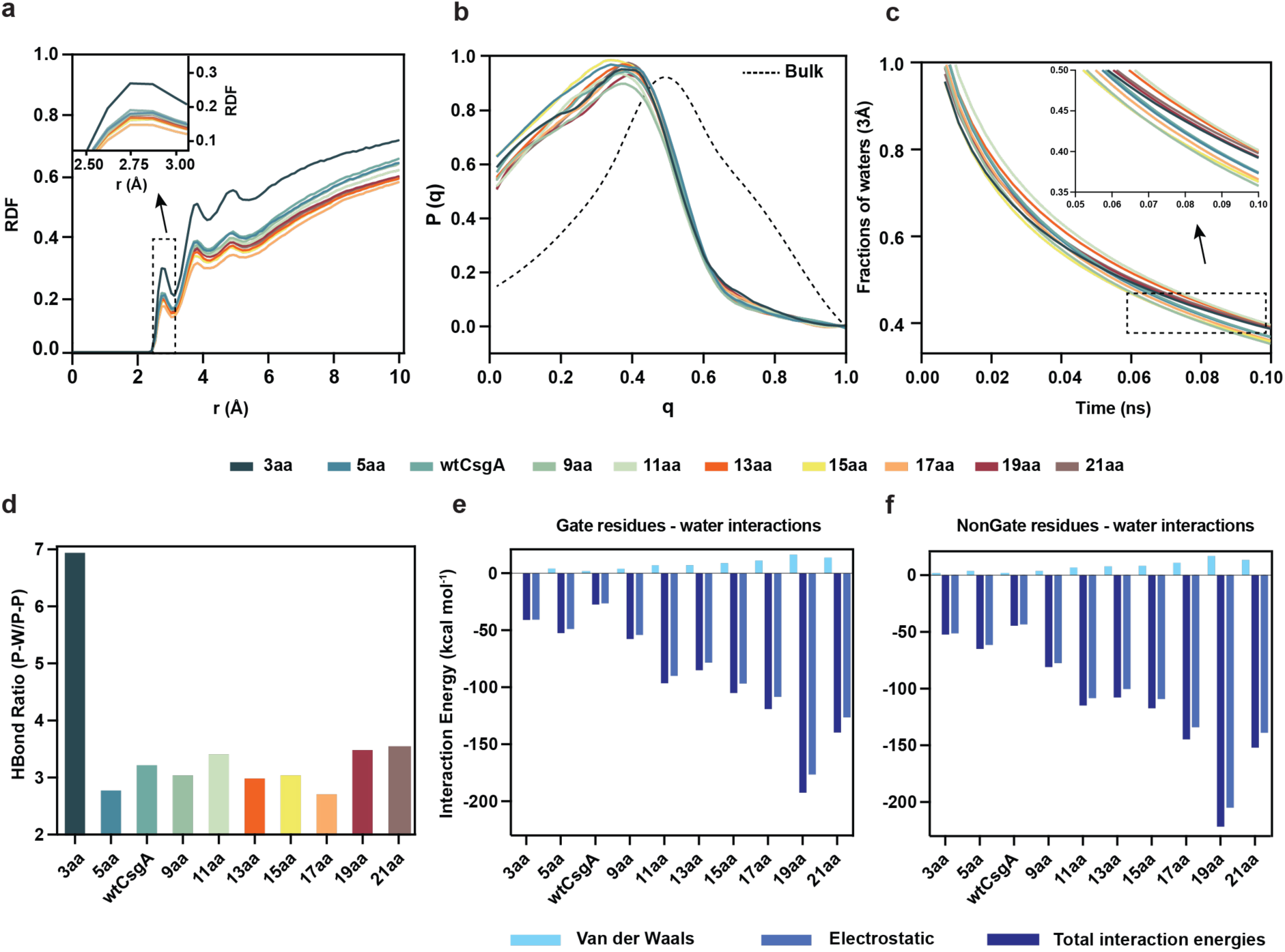
Mapping the Surface Hydration Landscape. **a.** Radial distribution function (RDF) between protein backbone heavy atoms and the oxygen atom of water as a function of radial distance. **b.** The orientational tetrahedral order (OTO) parameter of water around protein, and for bulk Water, is shown for comparison in a gray dotted line. **c.** The water-retention correlation among protein variants is shown in the inset, which is a zoomed-in view. **d.** The ratio of the hydrogen bonds between protein backbone and water oxygen atoms for all the variants. The bar plot shows the VdW, electrostatic, and total interaction energies between **e.** gate residues and **f.** non-gate residues with water. These energies are normalized per residue.

The tetrahedral order of waters is quantified by the orientational order parameter and shown in **Figure 4b, SI Appendix Fig. S13, and Table S5**. The peak shifts slightly for the 3aa, 11aa, and 15aa variants while remaining nearly constant for other variants. For all variants, the water order parameter peaks at approximately 0.4, lower than the 0.5 observed in bulk water, indicating distinct water ordering near the protein surface. **Figure 4c** illustrates the time evolution of the fraction of water molecules located within 3 Å of the protein surface for each variant. All variants show an initial rapid decay within the first nanosecond, reflecting the quick release or rearrangement of loosely bound water molecules. The inset highlights that CsgA, 3aa, and 5aa variants retain a higher fraction of interfacial water over time compared to other variants. This enhanced water retention may be attributed to the slightly higher percentage of polar residues present in these variants (**SI Appendix Fig. S14, SI Appendix Table S2**), which likely promotes stronger and more stable interactions with surrounding water molecules.

The electrostatic interactions of KR and non-KR with water were calculated in SI Appendix Table S6 and presented as bar plots on panels in **Figure 4e-f**. The van der Waals energies remained within a similar range (∼10 kcal mol^-1^) across all variants, whereas the electrostatic energy was more favorable for 19aa. The water interaction energies are comparable to those reported for the β-hairpin protein 2GB1 in the literature. (28) For non-key residue-water interactions, an increasing trend in electrostatic energies is observed with increasing variant size. To dissect these contributions, the interactions were separately tabulated for polar, nonpolar, and charged residues (**SI Appendix Table S7**, **SI Appendix Fig. S15**). The data confirm that electrostatic interactions involving polar and charged residues were more favorable than those involving nonpolar amino acids. In case of longer variants such as 15aa to 21aa, well-organized charged-residue patches (**SI Appendix Fig. S10**) are likely the primary factor contributing to the favorable residue-to-water electrostatic energy in **Figure 4f**.

### Biomanufacturing of Engineered β-Solenoid Variants and de novo autogenic ELM

Engineered CsgA variants with the highest confidence structural predictions from AlphaFold2 were selected for experimental validation (**Figure 2b**). Next, we aimed to determine whether the SECRETE platform(2), implemented in engineered *E. coli*, could successfully secrete, produce, and support self-assembly of these CsgA-inspired variants into extracellular β-solenoid structures. The amyloid cross-β structure of CsgA and engineered variants (3aa, 5aa, 9aa, 11aa, 13aa, 15aa, 17aa, 19aa, and 21aa) was assessed using a Congo Red binding assay (**Figure 5a**), which revealed robust dye binding across all variants comparable to CsgA. Minor variations in binding intensity likely reflect differences in amino acid composition, engineering-induced structural modifications, or variant-specific production levels. Wide-angle X-ray scattering (WAXS) analysis further confirmed the characteristic cross-β architecture, with inter-*β*-sheet and inter-*β*-strand *d*-spacing of approximately 0.98 nm and 0.46 nm, respectively, consistent with CsgA (**Figure 5b and 5c**). Finally, field-emission scanning electron microscopy (FESEM) revealed that the engineered CsgA variants self-assembled into extracellular nanofibers, confirming preservation of curli-like fibrillar morphology (**Figure 5d**).

**Figure 5.**
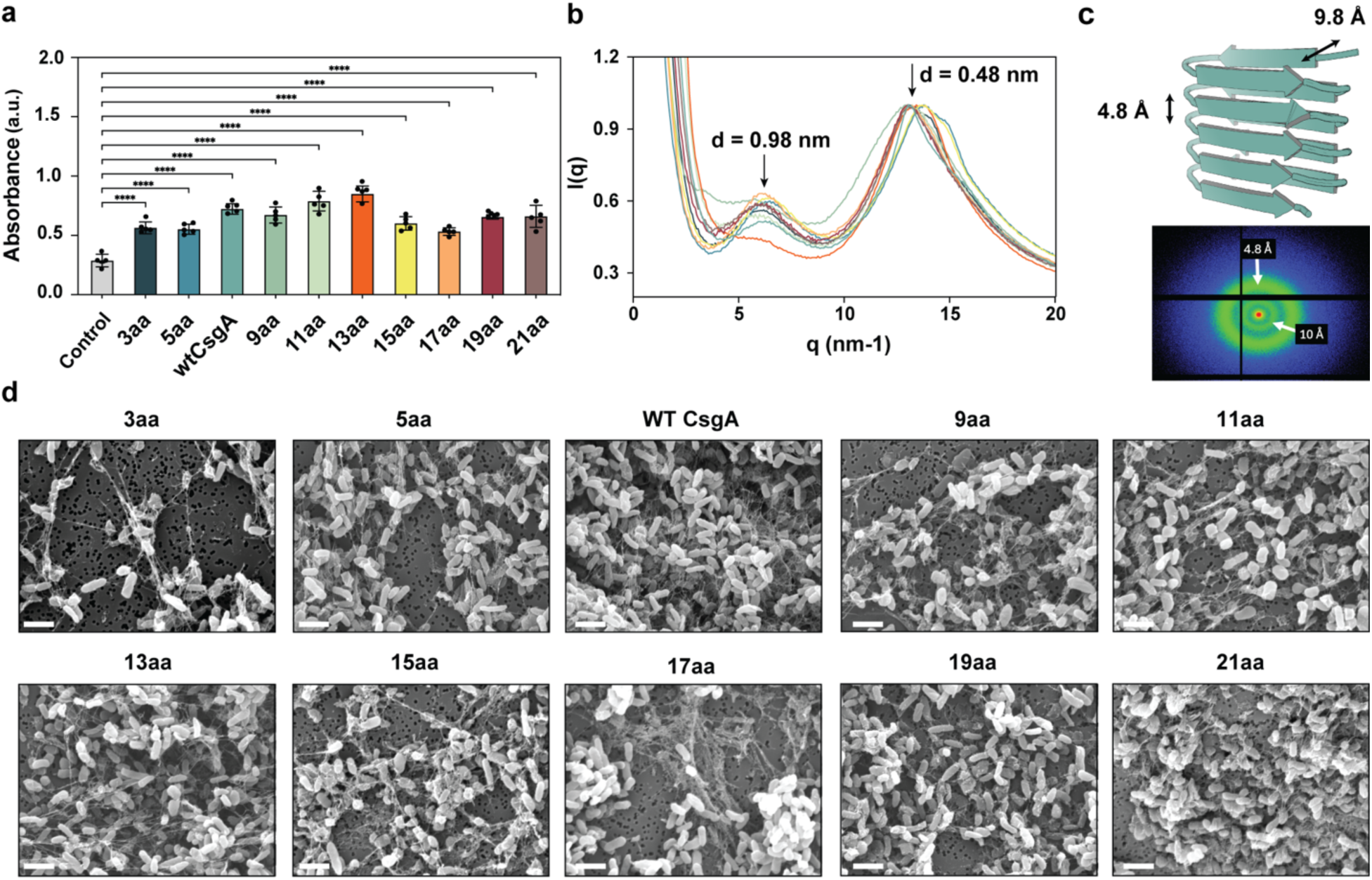
SECRETE platform for the production of engineered β-solenoid proteins by adopting the curli secretion machinery of *E. coli* (SECRETE). **a.** Congo red assay to determine the production of β-solenoid proteins by engineered *E. coli* and self-assembly into nanofibers, **b.** Wide-angle X-ray scattering (WAXS) analysis revealed β-cross characteristic of β-solenoid proteins, **c.** d-spacing (interplanar spacing) values of 0.98 nm and 0.46 nm, corresponding to inter-β-sheet and inter-β-strand distances, **d.** field-emission scanning electron microscopy (FESEM) images showed the protein nanofibers self-assembled in the extracellular milieu. Scale bar 1 µm.

To demonstrate the biomanufacturing of macroscopic engineered living materials (ELMs) from a library of CsgA β-solenoid variants with systematically tuned horizontal dimensions, we fabricated MECHS films (**Figure 6a**) from cultured bacterial biomass containing engineered curli protein nanofibers (40% w/w). To test this hypothesis, we performed tensile testing to quantify Young’s modulus, ultimate tensile strength, and elongation at break (**Figure 6b-e**). Deletion-based variants exhibited pronounced deviations from CsgA. Among these, the 3aa variant displayed the lowest Young’s modulus (14.44 ± 3.39 MPa), whereas the 5aa variant showed the highest stiffness within the designed library (44.48 ± 7.15 MPa). A similar trend was observed for ultimate tensile strength: the 3aa variant exhibited the lowest value (0.866 ± 0.106 MPa), while the 5aa variant showed one of the highest values (1.752 ± 0.161 MPa). In contrast, elongation at break followed an inverse trend, with the 3aa variant exhibiting the highest extensibility (63.09 ± 8.08%), and the 5aa variant one of the lowest (26.94 ± 2.6%). Insertion-based variants generally exhibited elongation at break values comparable to CsgA (50.22 ± 6.48%), with the exception of the 9aa (36.99 ± 6.05%) and 21aa (22.76 ± 3.94%) variants, which showed reduced extensibility. For Young’s modulus, most insertion-based variants exhibited lower stiffness than CsgA (31.78 ± 4.91 MPa), ranging from a minimum for the 17aa variant (17.58 ± 4.85 MPa) to a maximum for the 9aa variant (43.02 ± 7.82 MPa). A similar trend was observed for ultimate tensile strength, with the lowest value measured for the 21aa variant (1.137 ± 0.059 MPa) and the highest for the 9aa variant (1.865 ± 0.092 MPa), compared to CsgA (1.568 ± 0.066 MPa).

**Figure 6.**
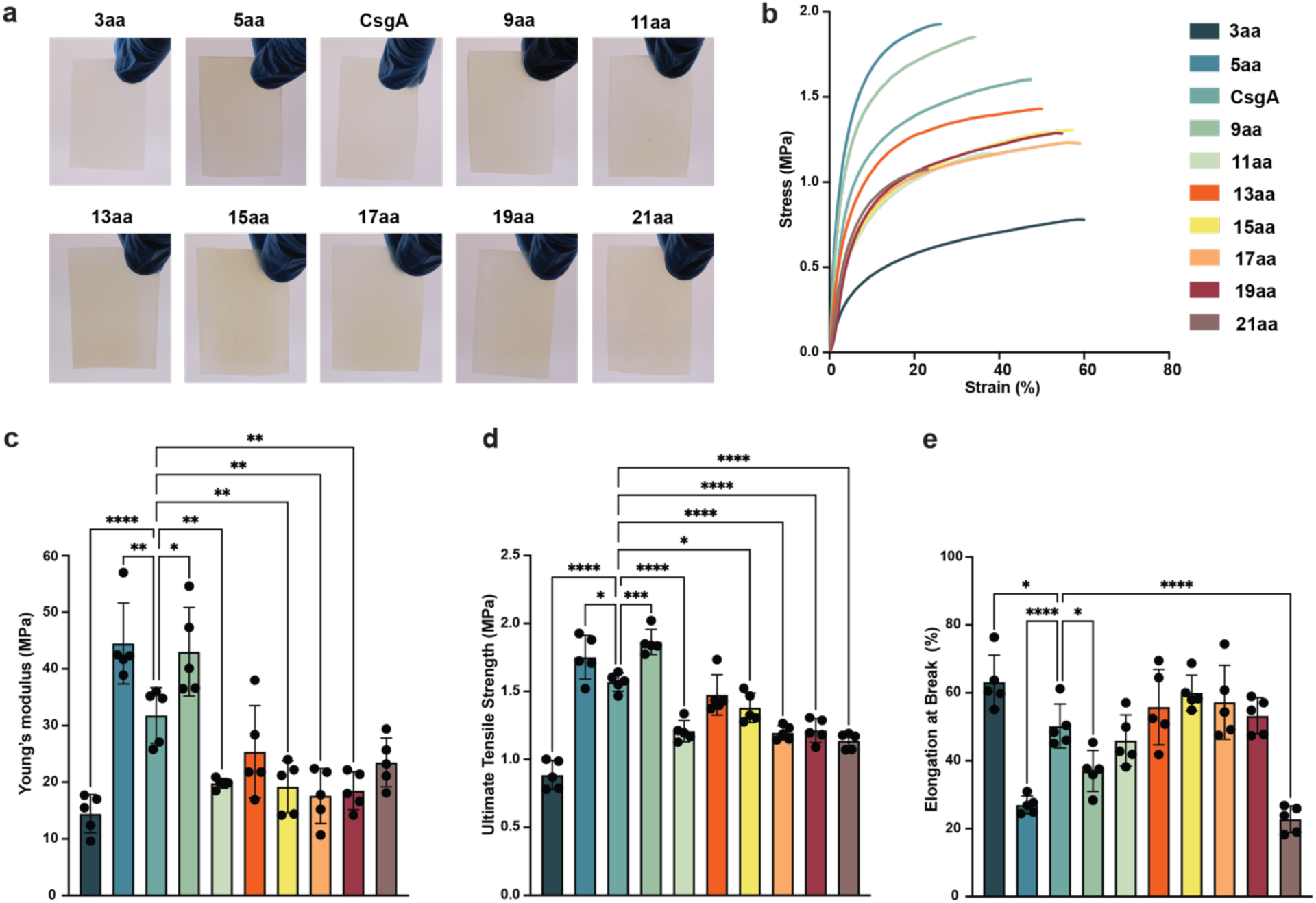
Biofabrication of MECHS film from the designed β-Solenoid variants. **a.** Representative photographs of MECHS designed variants, **b.** Representative stress-strain curves of MECHS films variants, **c.** Young’s modulus, **d.** Ultimate tensile strength and **e.** Elongation at the break. Biological replicates n = 5 for all β-Solenoid variants. Data represented as mean ± standard deviation for **c.** *p = 0.0145, **p = 0.0081, ****p < 0.0001; for **d.** *p = 0.0457, ***p = 0.0003; for **e.** *p = 0.0488, ****p < 0.0001. One-way ANOVA followed by Tukey’s multiple comparisons test.

## Conclusion

Building on evolutionary diversification of CsgA-like *β*-solenoid proteins, which primarily expands architecture along the vertical fiber axis, this work introduces an orthogonal design strategy that targets the horizontal dimension of the *β*-solenoid core. By rationally modulating β-strand length, we demonstrate a previously unexplored route for tuning β-solenoid architecture. Specifically, we engineered the horizontal dimension of the CsgA β-solenoid by varying the number of amino acids per β-strand, extending strand length from the native seven residues to variants spanning 3–21 residues. AI-based structural predictions using AlphaFold2 confirmed that this orthogonal design strategy preserves the β-solenoid fold across a broad range of strand lengths, supporting its feasibility as a general approach for tuning curli-based ECM architectures. Additionally, all-atom molecular dynamics simulations demonstrate that the structural stability of CsgA β-solenoid variants is governed by β-strand length, residue composition, and solvent interactions. The shortest variant (3aa) is unstable in aqueous environments, defining a lower bound for curli-like amyloid formation, whereas the 5aa variant exhibits exceptional stability in both aqueous and vacuum conditions, consistent with an optimal balance of hydrophobic and hydrophilic interactions. Variants with longer β-strands generally maintain structural integrity but occasionally undergo conformational rearrangements, and the distribution of polar and charged residues strongly influences their hydration behavior. Hydration and interaction analyses further highlight the stabilizing roles of key residues and solvent-mediated interactions, collectively linking molecular-level design features to emergent β-solenoid protein stability and material properties across environmental conditions. Importantly, we show that these engineered CsgA variants, including the less stable 3aa, can be produced by engineered *E. coli* using the SECRETE system, indicating that substantial modifications to the β-solenoid core remain compatible with native curli secretion, assembly, and autogenic ELM formation.

Importantly, the library of CsgA β-solenoid variants directly translates into emergent mechanical behavior at the macroscopic material scale. MECHS films fabricated from deletion-based variants exhibited differences in stiffness, strength, and extensibility relative to wild-type CsgA. The unstable 3aa variant exhibits the lowest Young’s modulus and tensile strength but the highest elongation at break, consistent with its reduced molecular stability and increased conformational flexibility. In contrast, the e 5aa variant produced the stiffest and strongest films within the designed library, while decreasing extensibility, in agreement with its enhanced β-solenoid stability predicted by simulations. Most insertion-based designs retained elongation at break values comparable to CsgA while exhibiting tunable stiffness and tensile strength as a function of strand length. Variants such as 9aa combined elevated stiffness and strength with reduced extensibility, whereas longer insertions (e.g., 17aa and 21aa) resulted in softer, weaker materials, suggesting that excessive horizontal expansion introduces local disorder or inefficient load transfer within the nanofiber network.

Structural plasticity of the CsgA β-solenoid is revealed by its ability to accommodate both deletion- and insertion-based variations in β-strand length, enabling controlled contraction or expansion of the horizontal core dimension while preserving the amyloid fold and self-assembly. Collectively, these findings position insertion-based β-strand elongation as a complementary strategy for fine-tuning mechanical performance without compromising assembly or material integrity. Such efforts could be integrated with rational design or directed evolution approaches, in combination with AI-based tools, to accelerate the design–build–test cycle for protein-based autogenic engineered living materials.

## Contributions

A.D.T. developed the initial concept. A.D.T. and A.M.B. streamlined the central idea and methodology. A.D.T. and H.H. designed experiments. H.H. performed AlphaFold2 structure prediction, H.H. and A.K. engineered *E. coli* to produce β-solenoid protein nanofibers, and S.S. performed molecular dynamics simulations and the analyses. E.J. performed WAXS analysis of β-solenoid proteins. A.M.B. performed SEM analysis. R.M. supervised E.C.; S.D. supervised S.S.; A.D.T. supervised H.H. The manuscript was drafted by H.H., S.S., and A.D.T. with input from all authors. All authors discussed the results and commented on the manuscript.

## Acknowledgment

Work was partly performed at the Advanced Research Computing Center, Nanoscale Characterization and Fabrication Laboratory (NCFL), and Institute for Critical Technology and Applied Science (ICTAS) at Virginia Tech. This work was supported by GlycoMIP, a National Science Foundation Materials Innovation Platform funded through Cooperative Agreement DMR-1933525.

## Materials and Methods

### Cell strains and plasmids

All experiments were carried out using the PQN4 strain of *E. coli*, derived from LSR10 (MC4100, ΔcsgA, λ(DE3), CamR) and lacking the curli operon (*ΔcsgBACEFG*). (29) The nine engineered *CsgA* amyloid domain genes were synthesized by Twist BioScience and cloned into the pET21d vector through overlap extension and isothermal Gibson Assembly (New England Biolabs). The complete plasmid sequences were verified using Plasmidsaurus. These plasmids also contained genes essential for curli biosynthesis, including *csgC*, *csgE*, *csgF*, and *csgG*. Ten distinct β-solenoid proteins—namely *E. coli CsgA*, 3aa, 5aa, 9aa, 11aa, 13aa, 15aa, 17aa, 19aa, and 21aa—were incorporated in place of the wild-type *E. coli CsgA*, following both the SEC (N-terminal signal sequence) and N22 (N-terminal curli-specific targeting sequence) to facilitate the secretion of *CsgA* fibers into the extracellular environment. Complete gene sequences, protein sequences, and primers utilized in this study are provided in **SI Appendix Fig. S7** and **Table S7**. Structure Prediction of Engineered Bacterial β-solenoid Proteins Using AlphaFold2

To predict the 3D structures of these 14 protein variants, we employed AlphaFold2 (version 2.1.1) (30) on the Tinkercliffs HPC cluster at the Advanced Research Computing Center, Virginia Tech, utilizing NVIDIA A100-80G architecture. The source code for AlphaFold2 is available at https://github.com/deepmind/alphafold. Structural images were generated using PyMOL (version 1.16). Batch protein structure predictions were performed, running for six recycles and generating five models per sequence. The models were ranked using the pLDDT score, with the highest-ranked model being used for subsequent analyses.

### Simulation setup

All-atom molecular dynamics (MD) simulations were conducted on wild-type CsgA and nine variants, featuring β-strands ranging from three (3aa) to twenty-one (21aa) amino acids, in an explicit aqueous environment using NAMD version 2.14.(31) The initial structural conformations for these β-solenoid proteins were derived from the AlphaFold2 server (30), based on experimentally determined FASTA sequences. Each amyloid structure was represented using the Charmm36m force field (updated July 2022).(32) The simulation box was solvated in a cubic simulation box with TIP3P (33) water, with 15 Å padding from the protein surface in all directions. To achieve charge neutrality, sodium ions were added to counterbalance the net charges of the amyloids, as detailed in Supplementary Table S1.

All the simulations were carried out with “rigidBonds all” that allowed a 2-fs integration timestep using the r-RESPA integrator (34) in the NPT ensemble, with periodic boundary conditions applied in all three directions. Non-bonded interactions were modeled using the standard 12-6 Lennard-Jones potential with a cutoff distance of 12 Å, while long-range electrostatics were computed via the Particle Mesh Ewald (PME) method. (35) Temperature and pressure were maintained at 300 K and 1 atm, respectively, using a Langevin thermostat (36) and a Parrinello-Rahman barostat.(37) Each system was first energy-minimized for 50,000 steps, followed by 500 ns MD simulations with trajectories recorded at 1 ps intervals. The first 250 ns were treated as equilibration, and the final 250 ns were used for structural analyses. Root-mean-square deviation (RMSD) calculations were performed over the full 500 ns to assess deviations from the initial AlphaFold2-predicted conformations, while all other structural, dynamical, and energetic analyses were conducted using only the production run.

### Analysis Method

The root-mean-square deviation (RMSD), hydrogen bond autocorrelation, radial distribution function (RDF), and Ramachandran plots were calculated utilizing the CPPTRAJ (38) and the MD Analysis library. (39, 40) Root-mean-square fluctuations (RMSF) of the amyloid structures and visualization were performed using the VMD software package. (41) For the hydrogen bond calculations, the HydrogenBondAnalysis (36) module of MDAnalysis was used, with a donor-acceptor distance cutoff of 3.0 Å and an angle cutoff of 30°. The secondary structure of the proteins was determined by the DSSP algorithm (42) using CPPTRAJ and MDTraj software. (43)

To characterize the local structure of liquid water in the vicinity of the amyloid proteins, the orientational tetrahedral order (OTO) parameter is calculated utilizing the Order software package. (44) For calculating the OTO parameter, an additional 1 ps simulation was carried out with a 1 fs timestep, taking the last trajectory snapshot of the 500 ns simulation for each of the variants. A custom code (available at https://github.com/Deshmukh-Group/Water-Residence-Persistence) is utilized to obtain the water retention time using MDAnalysis to track water molecules within 3.0 Å of protein backbone atoms across the last 50 ns trajectory. Retention time of water molecules is computed by assessing the fraction of common water molecules between frames separated by varying time intervals (Δt) and is normalized by the initial frame pairs, using parallel processing for efficiency. The resulting retention correlation decay is visualized as a function of Δt.

To facilitate efficient and parallel computation of residue-residue and residue-water interaction energies, an automated tool was developed leveraging the NAMD executable (available at https://github.com/Deshmukh-Group/ResidueResidue_Interaction_Energies_NAMD and https://github.com/Deshmukh-Group/ResidueWater_Interaction_Energies_NAMD). These interaction energies, both electrostatic and van der Waals contributions, were derived from non-bonded interactions within the 12 Å cutoff. For the residue-residue energy calculations, water molecules and ions were excluded from the system, and the final 50 ns of the simulation trajectories were analyzed with a stride of 100 ps. A filtering criterion was implemented, retaining only residue pairs with a center-of-mass separation of less than 12 Å in at least 60% of the trajectory frames for inclusion in the non-bonded energy calculations. The accuracy of these energy computations was cross-validated using the gRINN software package (45) and the VMD’s interaction energy calculation module. (41)

### Microbial production of engineered β-solenoid proteins

All plasmids, including CsgA, 3aa, 5aa, 9aa, 11aa,13aa, 15aa, 17aa,19aa, and 21aa, were introduced into the *E. coli* strain PQN4 via transformation. The transformed cells were streaked onto lysogeny broth (LB) agar plates supplemented with 100 µg/ml carbenicillin and 0.5% (w/v) glucose to repress T7RNAP through catabolite repression, followed by overnight incubation at 37 °C. A single colony from each transformation plate was selected and cultured separately in 5 ml of LB medium containing 100 µg/ml carbenicillin and 2% (w/v) glucose at 37 °C. These overnight cultures, designated as PQN4_ *CsgA*, PQN4_3aa, PQN4_5aa, PQN4_9aa, PQN4_11aa, PQN4_13aa, PQN4_15aa, PQN4_17aa, PQN4_19aa, and PQN4_21aa, were subsequently transferred to fresh 500 ml LB medium supplemented with 100 µg/ml carbenicillin. The cultures were incubated at 37 °C in a shaking incubator (225 rpm) for 48 hours to facilitate the expression of β-solenoid proteins and their assembly into functional amyloid nanofibers. As a negative control, PQN4 was transformed with an empty pET21d plasmid lacking all curli genes, including *csgA*.

### Quantitative Congo red dye binding assay

A 1 ml sample of the 48-hour, 500 ml bacterial culture was collected and centrifuged at 4000 rpm for 10 minutes. The resulting cell pellet was then resuspended in a 0.025 mM congo red solution prepared in phosphate-buffered saline (PBS) and incubated for 10 minutes. Following incubation, the cells were pelleted again by centrifugation at 14,000 rpm for 10 minutes. To assess amyloid nanofiber production, 200 µl of the supernatant was collected, and its absorbance at 490 nm was measured using a microplate reader. This absorbance was subtracted from that of the 0.025 mM Congo red solution in PBS and subsequently normalized by the OD600 value of the initial bacterial culture. The assay was conducted in four biological replicates.

### Field-emission scanning electron microscopy sample preparation and imaging

100 µl of the *E. coli* cultures producing protein variants were filtered onto polycarbonate membranes with a 0.22 μm pore size under vacuum and then placed in a fixative solution containing 1 ml of 4% glutaraldehyde and 1 ml of 4% paraformaldehyde buffer for 2 hours at room temperature. After fixation, the membranes were gently rinsed with water and subjected to a series of 200-proof ethanol washes at 25%, 50%, 75%, 100%, and 100% (v/v-1), each for 15 minutes. The samples were then processed using a critical point dryer. The dried membranes were mounted on Scanning Electron Microscopy sample holders using carbon adhesive and coated with a 10-20 nm Pt/Pd layer. Images were captured using a JEOL IT500 SEM equipped with a field-emission gun operating at 5–10 kV.

### Wide-angle X-ray scattering

Wide-angle X-ray scattering (WAXS) experiments on the aquaplastics were conducted using a Xenocs Xeuss 3.0 SAXS/WAXS system, equipped with a GeniX 3D Cu HFVLF microfocus X-ray source emitting Cu Kα radiation (λ = 0.154 nm). The sample-to-detector distance was set at 42 mm, and the *q*-range was calibrated using a Lanthanum hexaboride standard. Two-dimensional scattering patterns were recorded with a Dectris EIGER 4M detector, with an exposure time of 30 minutes. Data processing and reduction were carried out using XSACT software provided by Xenocs.

### Biofabrication of MECHS films

The 48 h cell cultures of PQN4_ CsgA, PQN4_3aa, PQN4_5aa, PQN4_9aa, PQN4_11aa, PQN4_13aa, PQN4_15aa, PQN4_17aa, PQN4_19aa, and PQN4_21aa were pelleted (5000 x *g*, 10 min) and washed with deionized water to remove residual culture media. 1g of wet biomass with engineered β-Solenoid nanofibers was dispersed in 5 ml of deionized water and 5 ml of 3% (w v^-1^) sodium dodecyl sulfate (gelator). After 2 h of incubation on a shaker at room temperature, the gelatinous biomass was developed and washed twice with 10 ml of deionized water. Next, to washed gelatinous biomass, 5 ml of 3% (w v^-1^) glycerol was added and mixed on a shaker for 1 h at room temperature. Finally, after incubation, the biomass was centrifuged and cast into a silicone mold and dried at 37 °C overnight.

### Statistics and reproducibility

All experiments described in this article were conducted at least three times (*n* ≥ 3) using biological replicates or distinct samples, as specified in the figure captions and relevant method sections. Data are reported as the mean with standard deviation. All data visualization and statistical analyses, including ordinary one-way ANOVA and Welch *t*-tests, were performed using GraphPad Prism 10. Additional data processing and reduction were conducted using XSACT software (Xenocs) for WAXS analysis. Microplate absorbance measurements were carried out using a BioTek Synergy H1 microplate reader (Agilent).

