## Supplementary Figures for "Rational Design Reveals Structural Plasticity of the CsgA β-Solenoid Enabling Programmable Autogenic Engineered Living Materials"

**Supplementary Table 1. Structure prediction of the b-solenoid proteins used in this study using AlphaFold2 based on pLDDT.**

- Very high (pLDDT > 90)
- High (90 > pLDDT > 70)
- Low (70 > pLDDT > 50)
- Very low (pLDDT < 50)

| | |
|---|---|
| **eCsgA-3aa** 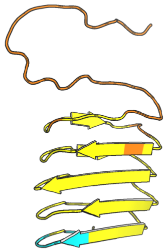 | 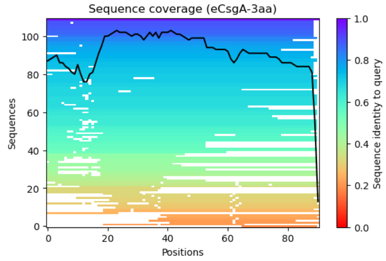  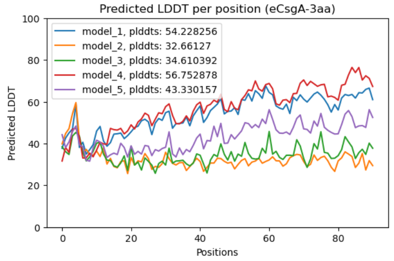  |
| **eCsgA-5aa** 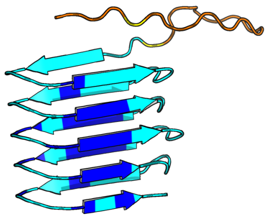 | 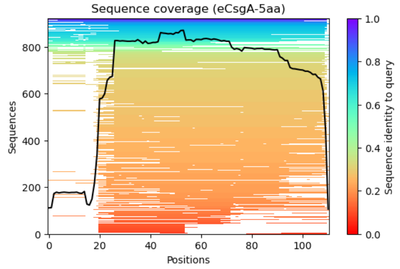  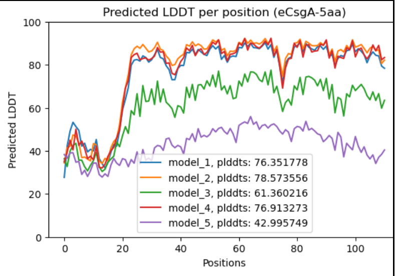  |
| ***E. coli* CsgA** 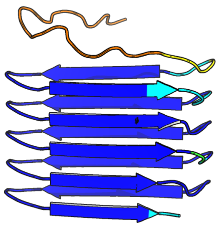 | 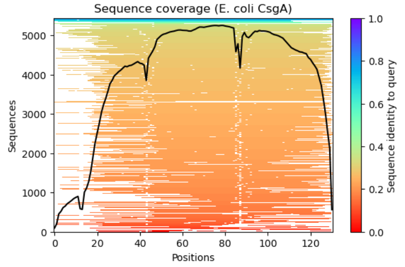  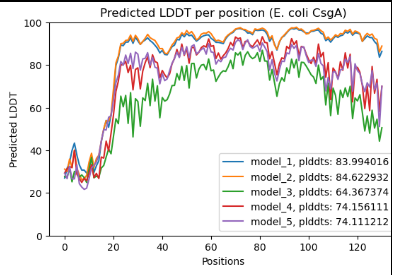  |

**CsgA-9aa**

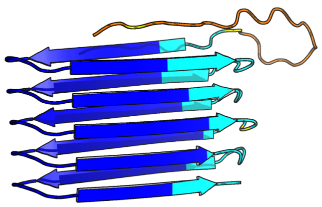

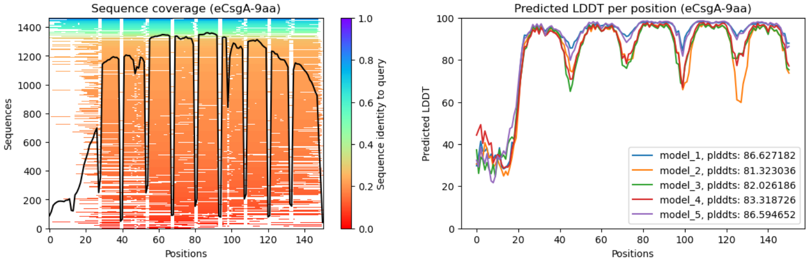

**CsgA-11aa**

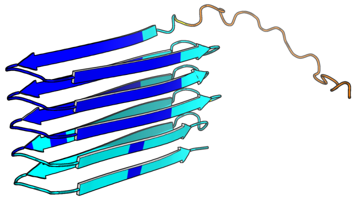

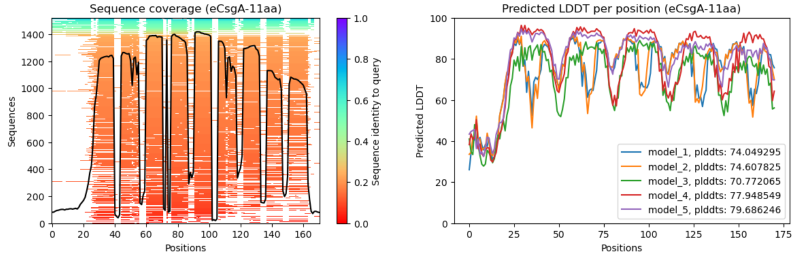

**CsgA-13aa**

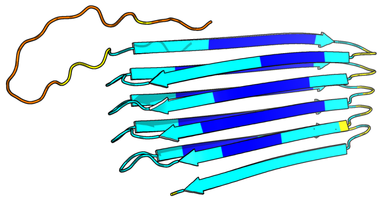

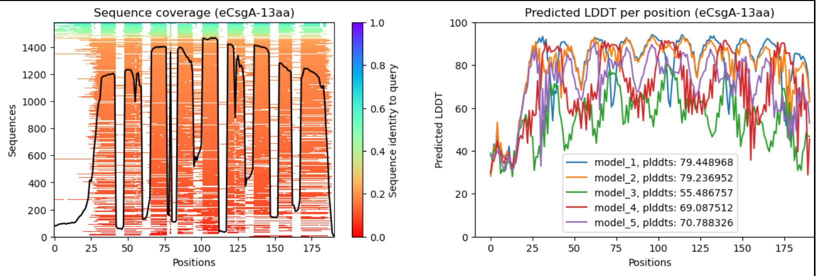

**CsgA-15aa**

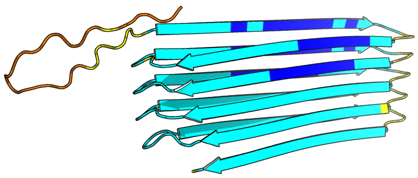

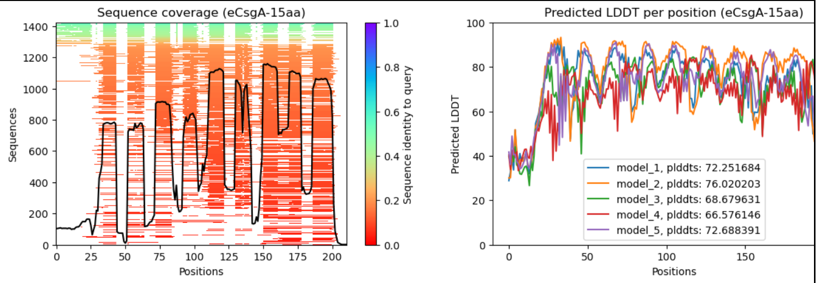

**CsgA-17aa**

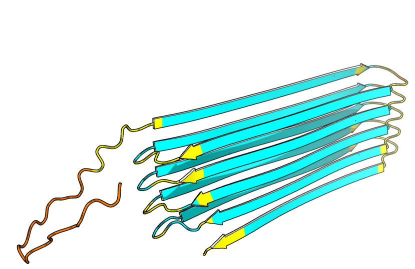

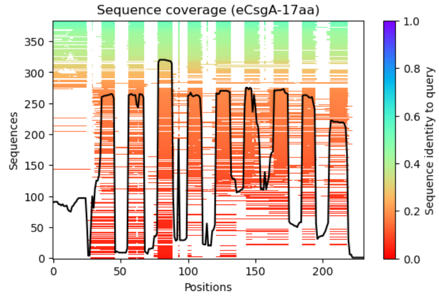

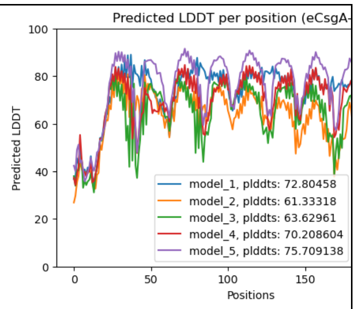

**CsgA-19aa**

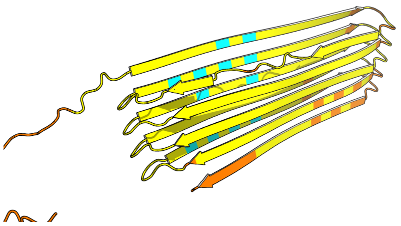

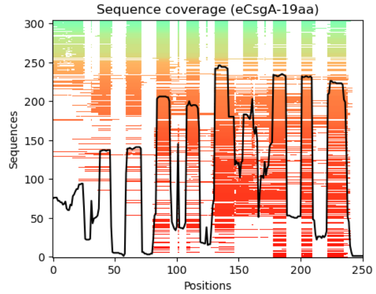

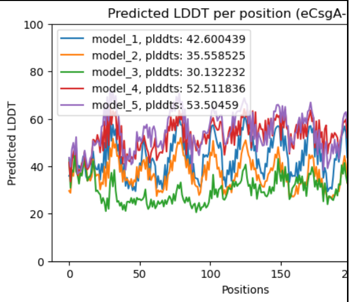

**CsgA-21aa**

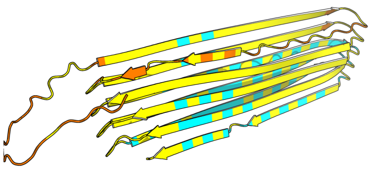

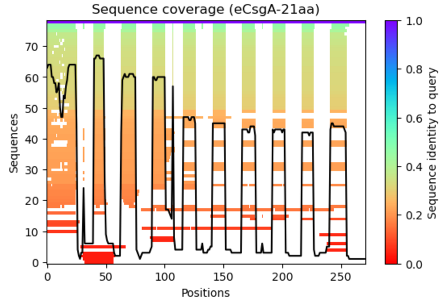

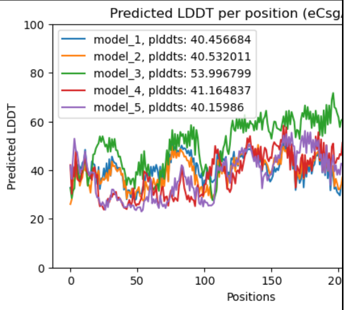

**Supplementary Figure S1**. Predicted structures of engineered CsgA β-solenoid protein variants using AlphaFold2 that were not used for experimental validation due to lower pLDDT scores.

**CsgA-5aa_2**

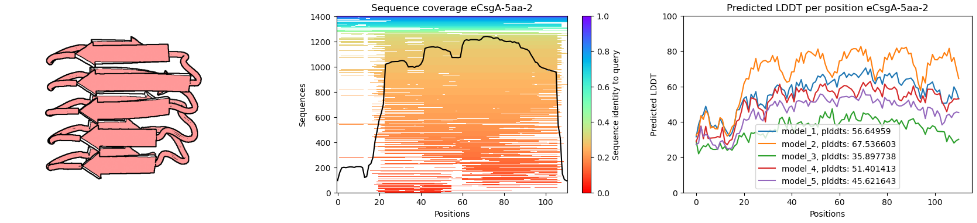

**CsgA-9aa_2**

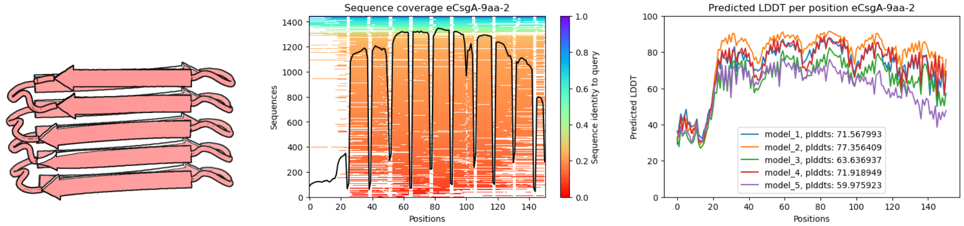

**CsgA-13aa_2**

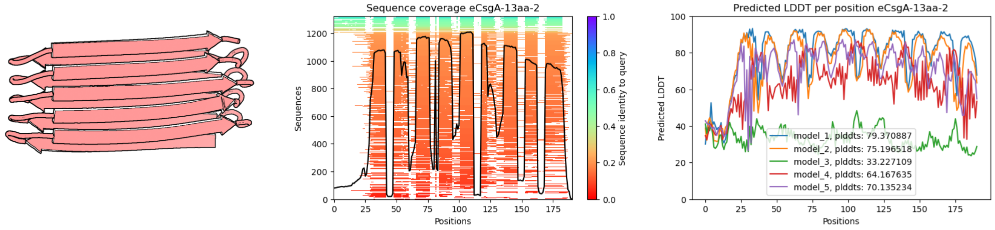

**CsgA-17aa_2**

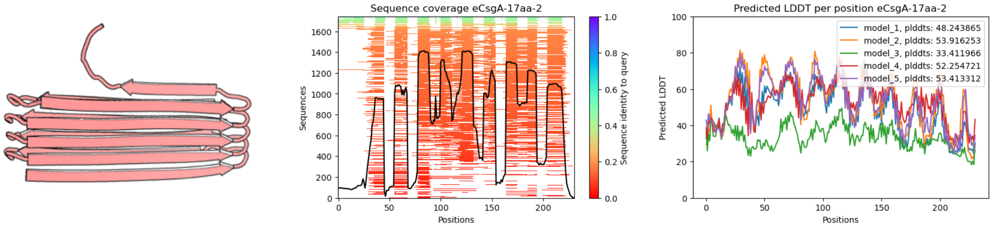

**CsgA-21aa_2**

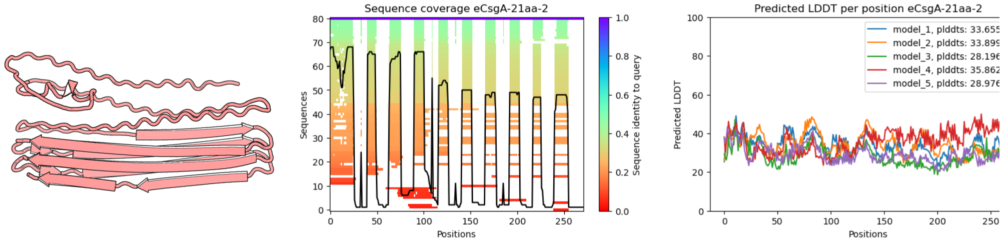

**Supplementary Figure S2.** 3D structure prediction of β-solenoid proteins based on Gravy hydrophobicity index.

| Hydrophilic | Hydrophobic |
|---|---|

**CsgA-3aa**

**GRAVY:** -1.387

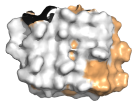

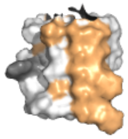

**CsgA-5aa**

**GRAVY:** -1.054

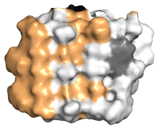

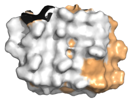

***E. coli*** **CsgA (wt CsgA)**

GRAVY: -0.718

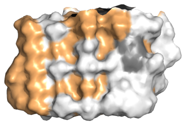

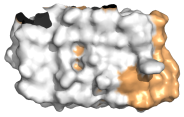

**CsgA-9aa**

**GRAVY:** -0.472

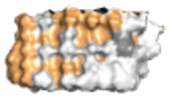

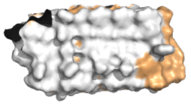

**CsgA-11aa**

**GRAVY:** -0.363

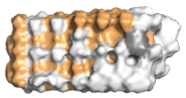

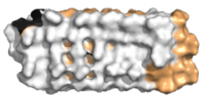

**CsgA-13aa**

GRAVY: -0.205

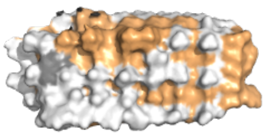

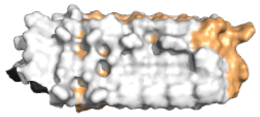

**CsgA-15aa**

**GRAVY:** -0.142

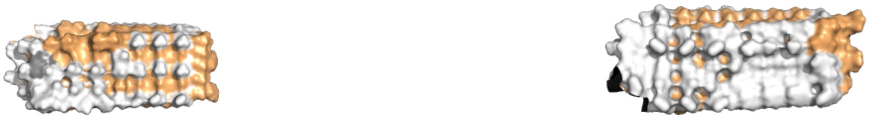

**CsgA-17aa**

**GRAVY:** -0.03

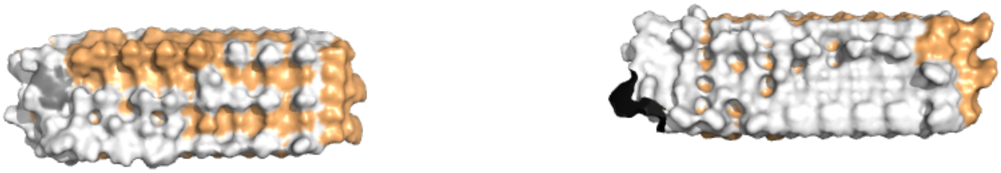

**CsgA-19aa**

**GRAVY:** 0.009

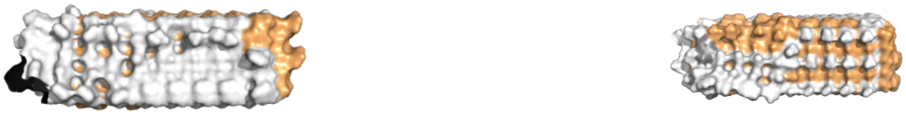

**CsgA-21aa**

**GRAVY:** 0.093

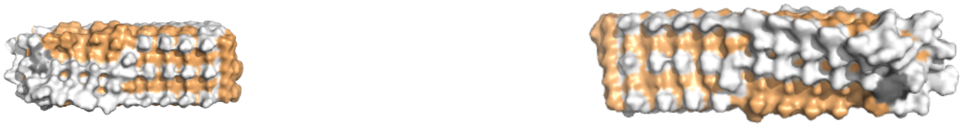

**Supplementary Figure 3.** Plasmid map of β-solenoid protein variants used in this study.

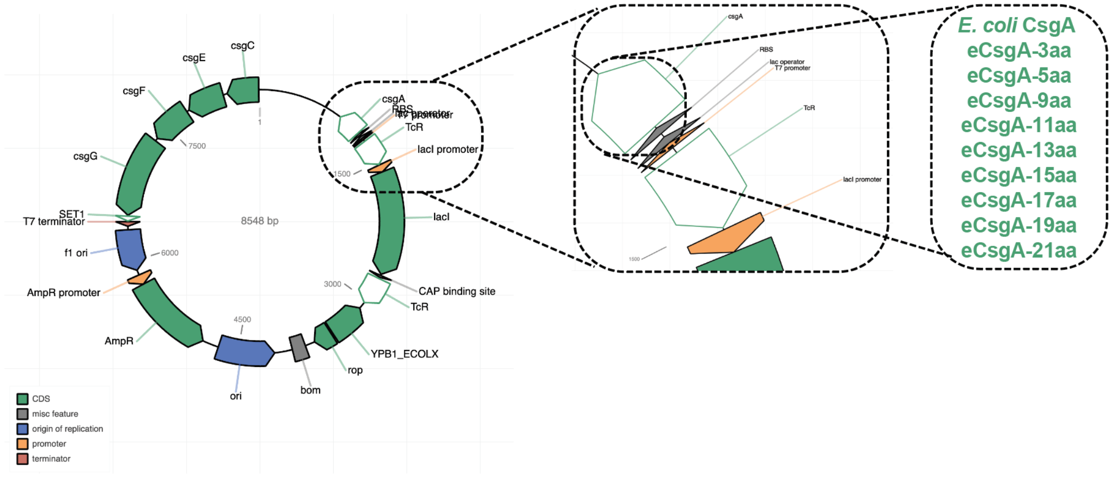

**Supplementary Table 2.** Sequences of engineered β-solenoid protein variants used in this study.

| Variant | Gene Sequence | Protein Sequence |
|---|---|---|
| **CsgA-3aa** | GGTGTTGTTCCTCAGTACGGCGGC<br>GGCGGTAACCACGGTGGTGGCGGT<br>AATAATAGCGGCCCAAATAGCAACC<br>AGTATGGCGGCGGGAACCTTCAAA<br>CCGACGCACGTAACAGCACACAGC<br>ATGGAGGTGGTAACGATCAGGGAA<br>GTGACGACTCGGATCAACGCGGGT<br>TCGGTAATACGCAGTGGAACGGCA<br>AGAATAGTACCCAGTTCGGTGGAG<br>GGAACGCGCAAACAGCATCCAATA<br>GTAACCAGGTAGGTTTCGGTAACAC<br>CCAATACTAA | GVVPQYGGGGNHGGGGNNSG<br>PNSNQYGGGNLQTDARNSTQH<br>GGGNDQGSDDSDQRGFGNTQ<br>WNGKNSTQFGGGNAQTASNS<br>NQVGFGNTQY* |
| **CsgA-5aa** | GGTGTTGTTCCTCAGTACGGCGGC<br>GGCGGTAACCACGGTGGTGGCGGT<br>AATAATAGCGGCCCAAATAGCGAAT<br>TGAACCAATACGGAGGCGGAAACA<br>GTGCATTACAGACTGATGCACGCAA<br>TTCTGATCTTACGCAGCACGGCGG<br>AGGTAACGGTGCCGATCAGGGCTC<br>GGACGACTCGTCTATCGACCAGCG<br>CGGATTTGGTAATAGCGCGACGCA<br>GTGGAATGGCAAAAACAGCGAAAT<br>GACACAGTTTGGAGGCGGGAACGG<br>AGCCGCGCAAACTGCCTCAAATTCC<br>TCCGTAAATCAGGTGGGTTTTGGTA<br>ACAACGCGACTCAATACTAA | GVVPQYGGGGNHGGGGNNSG<br>PNSELNQYGGGNSALQTDARN<br>SDLTQHGGGNGADQGSDDSSI<br>DQRGFGNSATQWNGKNSEMT<br>QFGGGNGAAQTASNSSVNQV<br>GFGNNATQY* |

| | | |
|---|---|---|
| ***E. coli* CsgA** | GGTGTTGTTCCTCAGTACGGCGGC<br>GGCGGTAACCACGGTGGTGGCGGT<br>AATAATAGCGGCCCAAATTCTGAGC<br>TGAACATTTACCAGTACGGTGGCG<br>GTAACTCTGCACTTGCTCTGCAAAC<br>TGATGCCCGTAACTCTGACTTGACT<br>ATTACCCAGCATGGCGGCGGTAAT<br>GGTGCAGATGTTGGTCAGGGCTCA<br>GATGACAGCTCAATCGATCTGACCC<br>AACGTGGCTTCGGTAACAGCGCTA<br>CTCTTGATCAGTGGAACGGCAAAAA<br>TTCTGAAATGACGGTTAAACAGTTC<br>GGTGGTGGCAACGGTGCTGCAGTT<br>GACCAGACTGCATCTAACTCCTCCG<br>TCAACGTGACTCAGGTTGGCTTTGG<br>TAACAACGCGACCGCTCATCAGTAC<br>TAA | GVVPQYGGGGNHGGGGNNSG<br>PNSELNIYQYGGGNSALALQTD<br>ARNSDLTITQHGGGNGADVGQ<br>GSDDSSIDLTQRGFGNSATLDQ<br>WNGKNSEMTVKQFGGGNGAA<br>VDQTASNSSVNVTQVGFGNNA<br>TAHQY* |
| **CsgA-9aa** | GGTGTTGTTCCTCAGTACGGCGGC<br>GGCGGTAATCATGGCGGGGGTGGA<br>AACAACTCGGGGCCGAACTCCGAG<br>TTGAATATATACATCTACCAATACG<br>GTGGTGGTAATAGTGCATTAGCCTT<br>GGCTCTTCAAACAGACGCCAGAAAT<br>TCTGACTTAACAATCACAATAACAC<br>AACACGGCGGTGGGAATGGTGCAG<br>ACGTCGGAGTTGGGCAAGGGAGTG<br>ACGACTCCTCAATCGACTTAACATT<br>AACACAACGTGGGTTCGGAAATTCG<br>GCCACTCTTGACTTGGACCAATGGA<br>ATGGTAAGAATTCCGAGATGACGGT<br>TAAGGTTAAGCAATTCGGCGGTGGT | GVVPQYGGGGNHGGGGNNSG<br>PNSELNIYIYQYGGGNSALALAL<br>QTDARNSDLTITITQHGGGNGA<br>DVGVGQGSDDSSIDLTLTQRGF<br>GNSATLDLDQWNGKNSEMTVK<br>VKQFGGGNGAAVDVDQTASNS<br>SVNVTVTQVGFGNNATAHAHQ<br>Y* |

| | | |
|---|---|---|
| | AATGGTGCCGCTGTCGACGTTGACCAAACTGCTTCTAATTCATCTGTCAATGTTACAGTTACACAAGTTGGTTTCGGTAATAATGCTACAGCTCACGCCCACCAATACTAA | |
| **CsgA-11aa** | GGTGTTGTTCCTCAGTACGGCGGCGGCGGTAATCATGGTGGTGGAGGCAACAACTCTGGTCCTAACAGTGAGTTGGAGTTAAATATATACATCTACCAATACGGCGGTGGGAATAGTGCTTCGGCCCTCGCATTAGCACTCCAAACGGACGCTCGTAATAGTGACCTTGACTTAACAATAACGATCACACAACACGGTGGTGGAAATGGTGCAGGTGCAGACGTAGGGGTCGGGCAAGGTTCTGACGACTCCAGTATAAGTATCGACTTGACGCTCACGCAACGGGGTTTCGGTAATTCCGCTTCGGCCACGTTGGACCTTGACCAATGGAATGGAAAGAATTCTGAGATGGAGATGACGGTCAAGGTAAAGCAATTCGGGGGGGGAAATGGAGCAGGTGCTGCAGTTGACGTAGACCAAACAGCCTCGAATTCTTCCGTATCTGTTAATGTTACGGTCACGCAAGTAGGGTTCGGAAATAATGCCAATGCCACAGCTCACGCTCACCAATACTAA | GVVPQYGGGGNHGGGGNNSGPNSELELNIYIYQYGGGNSASALALALQTDARNSDLDLTITITQHGGGNGAGADVGVGQGSDDSSISIDLTLTQRGFGNSASATLDLDQWNGKNSEMEMTVKVKQFGGGNGAGAAVDVDQTASNSSVSVNVTVTQVGFGNNANATAHAHQY* |
| **CsgA-13aa** | GGTGTTGTTCCTCAGTACGGCGGCGGCGGTAATCATGGTGGTGGTGGCAACAACTCAGGGCCCAACAGTGAGCTTGAGCTTAATATATACATATACAT | GVVPQYGGGGNHGGGGNNSGPNSELELNIYIYIYQYGGGNSASALALALALQTDARNSDLDLTITITITQHGGGNGAGADVGVGVGQ |

| | | |
|---|---|---|
| | CTACCAATACGGTGGTGGGAATAGT GCCTCGGCTTTAGCTTTAGCACTCG CATTGCAAACGGACGCACGGAATT CAGACCTTGACCTCACGATCACAAT CACAATCACACAACACGGTGGCGG AAATGGTGCAGGAGCAGACGTTGG TGTTGGTGTCGGTCAAGGTTCAGAC GACTCCTCGATATCCATCGACCTTA CACTCACTCTCACACAACGAGGGTT CGGTAATAGTGCTTCTGCAACTTTG GACTTAGACCTTGACCAATGGAATG GTAAGAATTCAGAGATGGAGATGAC TGTTAAGGTTAAGGTTAAGCAATTC GGAGGAGGAAATGGGGCAGGTGCT GCTGTAGACGTTGACGTTGACCAAA CTGCTAGTAATTCTTCGGTTTCAGT TAATGTCACAGTTACTGTAACGCAA GTCGGTTTCGGTAATAATGCCAATG CAACGGCTCACGCTCACGCACACC AATACTAA | GSDDSSISIDLTLTLTQRGFGNS ASATLDLDLDQWNGKNSEMEM TVKVKVKQFGGGNGAGAAVDV DVDQTASNSSVSVNVTVTVTQ VGFGNNANATAHAHAHQY* |
| **CsgA-15aa** | GGTGTTGTTCCTCAGTACGGCGGC GGCGGTAATCATGGTGGTGGCGGC AACAACTCGGGGCCCAACTCGGAG TTGGAGCTCGAGTTGAATATATACA TATACATATACCAATACGGAGGAGG AAATTCCGCTTCTGCATCCGCACTT GCACTTGCATTAGCTTTGCAAACGG ACGCTCGTAATTCTGACCTTGACTT AGACTTGACGATAACTATAACGATC ACGCAACACGGTGGTGGTAATGGT GCTGGTGCCGGAGCAGACGTAGGG | GVVPQYGGGGNHGGGGNNSG PNSELELELNIYIYIYQYGGGNS ASASALALALALQTDARNSDLD LDLTITITITQHGGGNGAGAGAD VGVGVGQGSDDSSISISIDLTLT LTQRGFGNSASASATLDLDLDQ WNGKNSEMEMEMTVKVKVKQ FGGGNGAGAGAAVDVDVDQT ASNSSVSVSVNVTVTVTQVGF GNNANANATAHAHAHQY* |

| | | |
|---|---|---|
| | GTAGGAGTTGGTCAAGGTTCCGAC<br>GACTCTAGTATAAGTATATCAATAG<br>ACTTAACGTTAACTCTTACACAAAG<br>AGGGTTCGGAAATTCAGCCTCTGCT<br>TCGGCCACTCTCGACTTGGACCTTG<br>ACCAATGGAATGGTAAGAATAGTGA<br>GATGGAGATGGAGATGACTGTCAA<br>GGTTAAGGTCAAGCAATTCGGCGG<br>TGGTAATGGAGCCGGTGCCGGTGC<br>CGCTGTAGACGTAGACGTTGACCA<br>AACTGCAAGTAATTCCTCAGTATCG<br>GTTTCAGTAAATGTCACAGTTACGG<br>TTACGCAAGTTGGTTTCGGGAATAA<br>TGCTAATGCTAATGCTACGGCCCAC<br>GCACACGCTCACCAATACTAA | |
| **CsgA-17aa** | GGTGTTGTTCCTCAGTACGGCGGC<br>GGCGGTAATCATGGGGGCGGTGG<br>GAACAACTCTGGTCCCAACTCAGAG<br>TTAGAGCTCGAGTTGAATATCTACA<br>TCTACATATACATATACCAATACGGT<br>GGCGGAAATAGTGCCTCAGCTAGT<br>GCATTGGCCCTTGCTTTAGCTCTCG<br>CATTACAAACGGACGCTCGTAATAG<br>TGACCTCGACTTAGACTTGACGATC<br>ACTATAACAATAACAATCACACAAC<br>ACGGTGGTGGGAATGGTGCAGGAG<br>CCGGGGCTGACGTAGGTGTCGGTG<br>TTGGGGTTGGGCAAGGTTCTGACG<br>ACAGTTCTATCTCCATCTCAATCGA<br>CCTTACTCTTACGCTCACTTTAACG<br>CAAAGAGGATTCGGAAATTCTGCAT | GVVPQYGGGGNHGGGGNNSG<br>PNSELELELNIYIYIYIYQYGGGN<br>SASASALALALALALQTDARNS<br>DLDLDLTITITITITQHGGGNGAG<br>AGADVGVGVGVGQGSDDSSISI<br>SIDLTLTLTLTQRGFGNSASASA<br>TLDLDLDLDQWNGKNSEMEME<br>MTVKVKVKVKQFGGGNGAGA<br>GAAVDVDVDVDQTASNSSVSV<br>SVNVTVTVTVTQVGFGNNANA<br>NATAHAHAHAHQY* |

| | | |
|---|---|---|
| | CCGCATCAGCCACTCTCGACCTTGA<br>CCTTGACTTGGACCAATGGAATGGA<br>AAGAATTCAGAGATGGAGATGGAG<br>ATGACGGTTAAGGTTAAGGTTAAGG<br>TTAAGCAATTCGGTGGTGGTAATGG<br>GGCAGGTGCCGGGGCCGCCGTTG<br>ACGTTGACGTTGACGTCGACCAAAC<br>AGCATCAAATAGTTCGGTTTCTGTT<br>TCAGTTAATGTTACTGTAACTGTAAC<br>GGTCACGCAAGTTGGTTTCGGAAAT<br>AATGCCAATGCTAATGCCACGGCC<br>CACGCACACGCCCACGCCCACCAA<br>TACTAA | |
| **CsgA-19aa** | GGTGTTGTTCCTCAGTACGGCGGC<br>GGCGGTAACCACGGTGGTGGCGGT<br>AATAATAGCGGCCCAAATAGCGAAT<br>TGGAGTTGGAATTAGAGCTTAACAT<br>TTATATCTACATCTATATCTACCAGT<br>ATGGAGGCGGTAATTCTGCATCAG<br>CATCGGCGTCAGCCCTTGCACTTG<br>CCCTTGCTTTGGCGCTGCAGACAG<br>ATGCGCGTAATTCAGATCTTGACCT<br>GGACCTTGACCTTACAATCACAATC<br>ACTATCACTATCACACAACACGGTG<br>GGGGTAATGGAGCCGGTGCTGGAG<br>CCGGTGCAGATGTAGGCGTGGGTG<br>TTGGTGTCGGGCAAGGGTCGGACG<br>ACTCTTCAATTTCAATTTCGATTAGC<br>ATTGACCTGACCCTGACTTTGACAC<br>TGACTCAACGCGGGTTCGGAAATTC<br>GGCGTCAGCGTCTGCATCCGCTAC | GVVPQYGGGGNHGGGGNNSG<br>PNSELELELELNIYIYIYIYQYGG<br>GNSASASASALALALALQTD<br>ARNSDLDLDLDLTITITITITQHG<br>GGNGAGAGAGADVGVGVGVG<br>QGSDDSSISISISIDLTLTLTLTQ<br>RGFGNSASASASATLDLDLDLD<br>QWNGKNSEMEMEMEMTVKVK<br>VKVKQFGGGNGAGAGAGAAV<br>DVDVDVDQTASNSSVSVSVSV<br>NVTVTVTVTQVGFGNNANANA<br>NATAHAHAHAHQY* |

| | | |
|---|---|---|
| | TTTGGACCTTGATCTGGATCTGGAT CAATGGAACGGCAAAAACTCTGAGA TGGAGATGGAGATGGAAATGACTG TAAAAGTGAAGGTGAAAGTCAAGCA GTTTGGTGGGGGAAATGGGGCCGG AGCTGGGGCCGGCGCAGCGGTCG ACGTGGACGTGGACGTTGACCAAA CAGCGAGTAATAGCAGCGTGAGTG TATCTGTATCTGTGAATGTGACGGT CACAGTAACAGTGACGCAAGTTGG CTTTGGTAATAATGCCAATGCAAAC GCAAATGCAACAGCGCATGCACAC GCCCATGCCCACCAGTACTAA | |
| **CsgA-21aa** | GGTGTTGTTCCTCAGTACGGCGGC GGCGGTAACCACGGTGGTGGCGGT AATAATAGCGGCCCAAATAGTGAGT TGGAGCTTGAGTTAGAGTTAAATAT CTACATCTACATATACATATACATCT ACCAATACGGTGGTGGTAATAGTGC CAGTGCTTCAGCAAGTGCATTAGCC TTAGCTTTGGCCTTGGCCTTAGCAT TACAAACGGACGCACGAAATTCAGA CTTGGACCTTGACTTAGACTTAACA ATAACGATCACAATAACGATAACTA TCACTCAACACGGTGGTGGGAATG GGGCAGGTGCCGGTGCAGGGGCC GACGTCGGAGTTGGGGTCGGAGTT GGTGTTGGGCAAGGTTCAGACGAC TCGAGTATCTCTATCTCCATCAGTA TCGACTTGACTTTGACTCTTACACTT ACTCTCACACAAAGAGGGTTCGGAA | GVVPQYGGGGNHGGGGNNSG PNSELELELELNIYIYIYIYIYQYG GGNSASASASALALALALAL QTDARNSDLDLDLDLTITITITITI TQHGGGNGAGAGAGADVGVG VGVGVGQGSDDSSISISISIDLT LTLTLTLTQRGFGNSASASASA TLDLDLDLDLDQWNGKNSEME MEMEMTVKVKVKVKVKQFGGG NGAGAGAGAAVDVDVDVDVD QTASNSSVSVSVSVNVTVTVTV TVTQVGFGNNANANANATAHA HAHAHAHQY* |

| | |
|---|---|
| | ATTCAGCATCTGCTTCTGCATCTGC<br>CACGTTAGACTTAGACCTCGACTTG<br>GACTTGGACCAATGGAATGGTAAGA<br>ATTCAGAGATGGAGATGGAGATGG<br>AGATGACTGTTAAGGTCAAGGTTAA<br>GGTTAAGGTTAAGCAATTCGGTGGT<br>GGTAATGGTGCTGGAGCCGGGGCC<br>GGAGCAGCTGTTGACGTCGACGTA<br>GACGTAGACGTTGACCAAACTGCTT<br>CCAATTCCAGTGTATCAGTCAGTGT<br>ATCGGTTAATGTTACGGTCACGGTC<br>ACGGTTACGGTCACGCAAGTTGGG<br>TTCGGAAATAATGCAAATGCTAATG<br>CCAATGCCACTGCACACGCACACG<br>CCCACGCCCACGCACACCAATACT<br>AG |

**Supplementary Figure** X: Snapshots of protein from the last 250 ns of MD simulations. All variants and WT except 3aa retain the core β-solenoid secondary structure. Elevated levels of fluctuation are noted in the N- and C-terminal regions of the proteins.

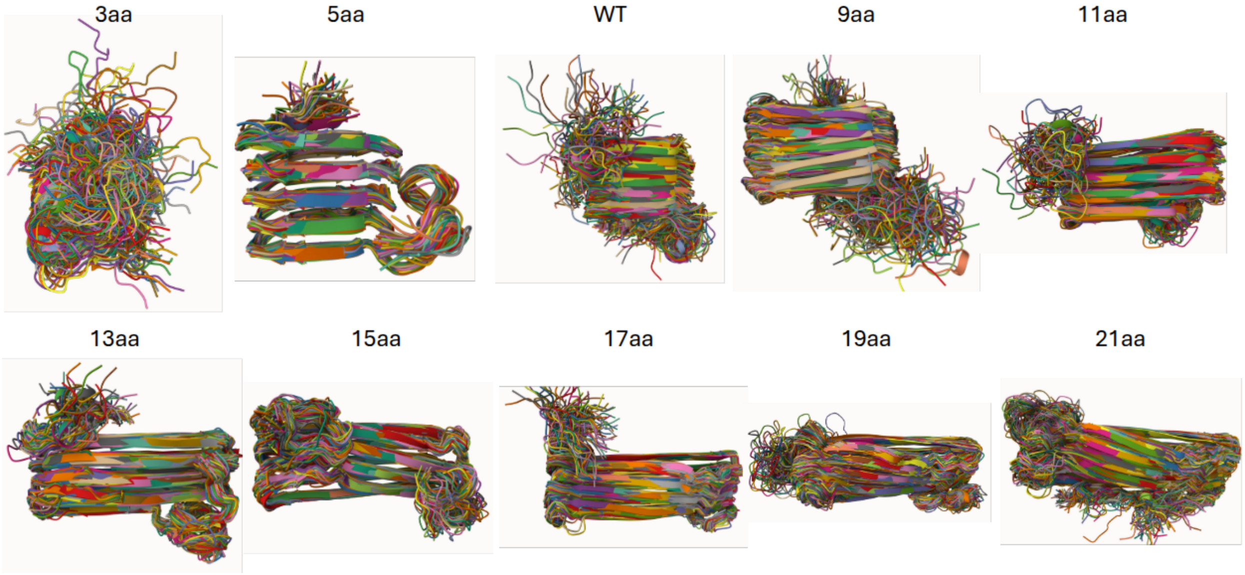

**Supplementary Table S1:** List of the total number of amino acid residues, net charge, percentage of positively and negatively charged residues, and total charged residue percentage for the CsgA WT and the nine variants.

| CsgA variants | Number of amino acid Residues | MW g/mol | GRAVY | Net Charge | Positively Charged Residues (%) | Negatively Charged Residues (%) | Total Charged Residues (%) |
|---|---|---|---|---|---|---|---|
| **WT** | 131 | 13092.69 | -0.718 | -6e | 3.05 | 7.63 | 10.69 |
| **3aa** | 91 | 9094.14 | -1.387 | -2e | 3.30 | 5.49 | 8.79 |
| **5aa** | 111 | 10969.17 | -1.054 | -5e | 2.70 | 7.21 | 9.91 |
| **9aa** | 151 | 15216.21 | -0.472 | -7e | 3.31 | 7.95 | 11.26 |
| **11aa** | 171 | 17091.25 | -0.363 | -10e | 2.92 | 8.77 | 11.70 |
| **13aa** | 191 | 19214.77 | -0.205 | -11e | 3.14 | 8.90 | 12.04 |
| **15aa** | 211 | 21089.81 | -0.142 | -14e | 2.84 | 9.48 | 12.32 |
| **17aa** | 231 | 23213.33 | -0.03 | -15e | 3.03 | 9.52 | 12.55 |
| **19aa** | 251 | 25088.37 | 0.009 | -18e | 2.79 | 9.96 | 12.75 |
| **21aa** | 271 | 27211.89 | 0.093 | -19e | 2.95 | 9.96 | 12.92 |

**Supplementary Table S2:** List of the total number of amino acid residues, polar (not including the charged residues), charged, and nonpolar residue percentages in the β-solenoid core (total residues – N22 terminal residues) for the CsgA WT and the nine variants.

| CsgA variants | β-solenoid core Residue | Polar Residue (%) | Charged Residue (%) | Non-Polar Residue (%) |
|---|---|---|---|---|
| **WT** | 109 | 43.12 | 12.84 | 44.04 |
| **3aa** | 69 | 53.62 | 11.59 | 34.78 |
| **5aa** | 89 | 47.19 | 12.36 | 40.45 |
| **9aa** | 129 | 40.31 | 13.18 | 46.51 |
| **11aa** | 149 | 38.26 | 13.42 | 48.32 |
| **13aa** | 169 | 36.69 | 13.61 | 49.70 |
| **15aa** | 189 | 35.45 | 13.76 | 50.79 |
| **17aa** | 209 | 34.45 | 13.88 | 51.67 |
| **19aa** | 229 | 33.62 | 13.97 | 52.40 |
| **21aa** | 249 | 32.93 | 14.06 | 53.01 |

**RMSF and Ramachandran analysis:** Root-mean-square fluctuation (RMSF) was calculated to assess residue-level local dynamics over a 5-frame interval during the final 50 ns of the simulation trajectory, as shown in Fig. S2a. For well-ordered β-strands, RMSF values remained consistently low, below 1.5 Å, indicating minimal conformational variability across all variants except 3AA. The 3AA variant exhibited elevated fluctuations in the core β-sheet regions, suggesting an unfolding tendency. The N-terminal regions (first 22 residues in each protein) displayed higher RMSF values, reflecting their flexible, coil-like conformations. Additionally, intermittent fluctuations in the C-terminal β-strand contributed to increased RMSF values for the 13AA, 15AA, and 19AA variants compared to the well-ordered core β-strands.

The Ramachandran plot, shown in Fig. S2b, displays characteristic signatures of parallel β-strands at approximately $\phi$ = -120° and $\psi$ = 120°, appearing as a distinct node (colored red) in the heatmap, indicative of a high β-sheet population for all variants except 3AA. In the 3AA variant, the β-sheet population is diminished, with increased population in the β-turn and coil regions suggesting structural changes and unfolding. Additionally, type II β-turns, which connect consecutive β-strands, were identified at approximately $\phi$ = -60° and $\psi$ = 125°. For all other variants, the β-sheet population showed minimal variation, indicating that their secondary structures remained stable in the simulation. In the case of the 3aa, due to the unfolding of the structure, the absence of β-strand signatures was noted, instead the β-turn regions being populated, corroborating the instability of the secondary β-sheet structure.

**Supplementary Figure S2:** (a) RMSF of the WT CsgA and the nine variants. (b) Ramachandran plot as a function of dihedral angles $\phi$ and $\psi$. The heatmaps show the relative concentration of the dihedral angles corresponding to the different secondary conformations in each of the cases.

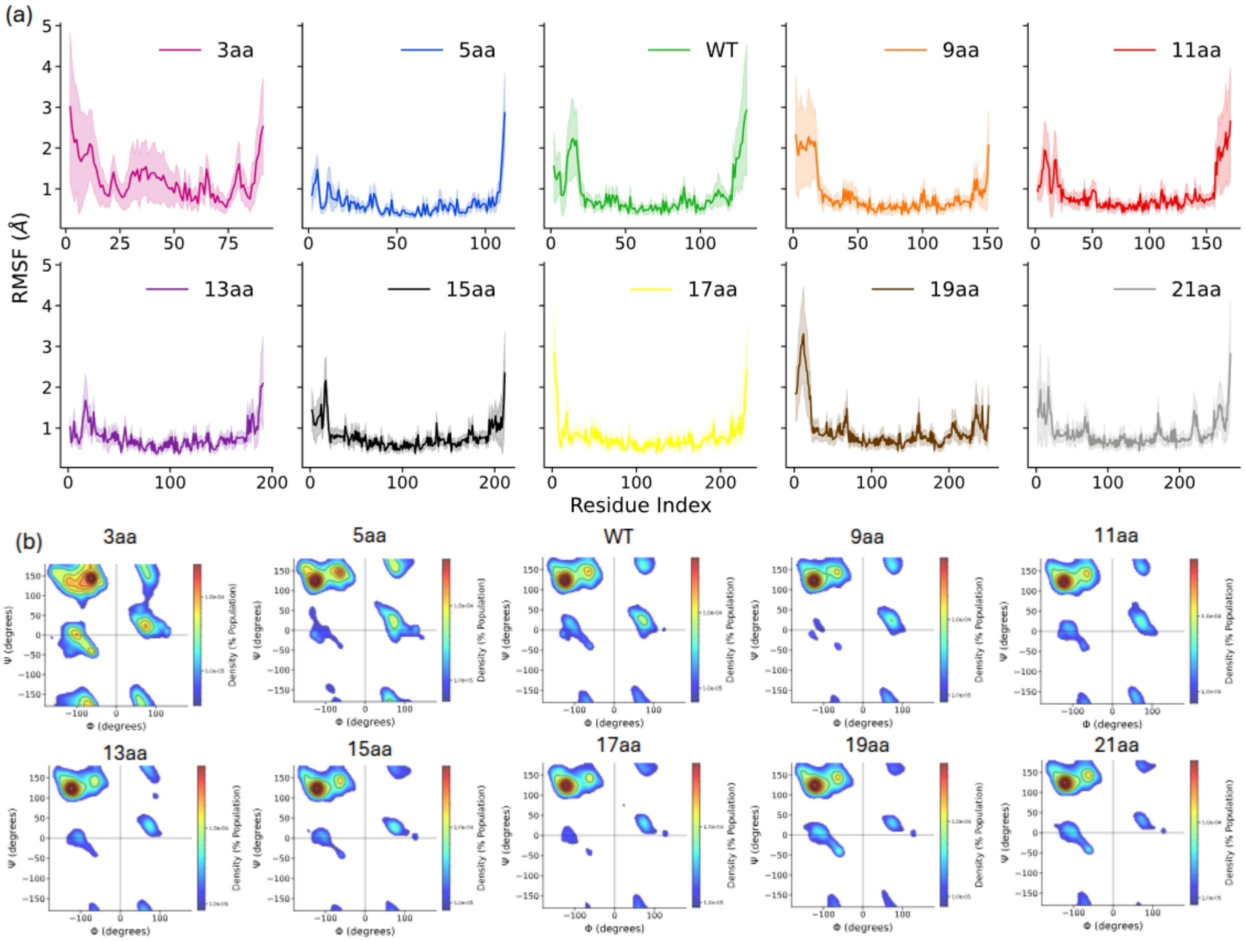

**Supplementary Figure S3:** (a) Comparison of RMSD for protein heavy atoms in aqueous (in color) and without water (in gray). (b) The radar plot shows the VdW, electrostatic, and total interaction energies measured between gate-to-gate residues, gate-to-all residues, and non-gate-to-all residues in the absence of water.

**Effect of water on CsgA variants' stability:** Water is known to influence proteins' secondary structure stability with favorable interactions with hydrophilic residues on the protein surface, while hydrophobic residues are typically sequestered within the protein core, shielded from water [13] . To evaluate the impact of water interaction on β-solenoid stability, simulations of these proteins were conducted separately in vacuum, with the resulting RMSD values of protein heavy atoms compared to those in aqueous conditions, as shown in Fig. S3 (a)-(b). For the 3aa variant, the secondary structure was found to remain stable in the absence of water, in contrast to its unfolding behavior in water, suggesting that water interactions are responsible for structural deformation in this case [14]. On the contrary, the 5aa variant was noted to exhibit exceptional stability in both aqueous and vacuum environments, with closely aligned RMSD values. For the other variants, the β-proteins were generally observed to be more stable in the absence of water, except for the 9aa, 15aa, and 19aa variants, where RMSD values were comparable between the two conditions. These results confirm that solvation affects β-solenoids to varying degrees, contingent upon the location and composition of amino acid residues.

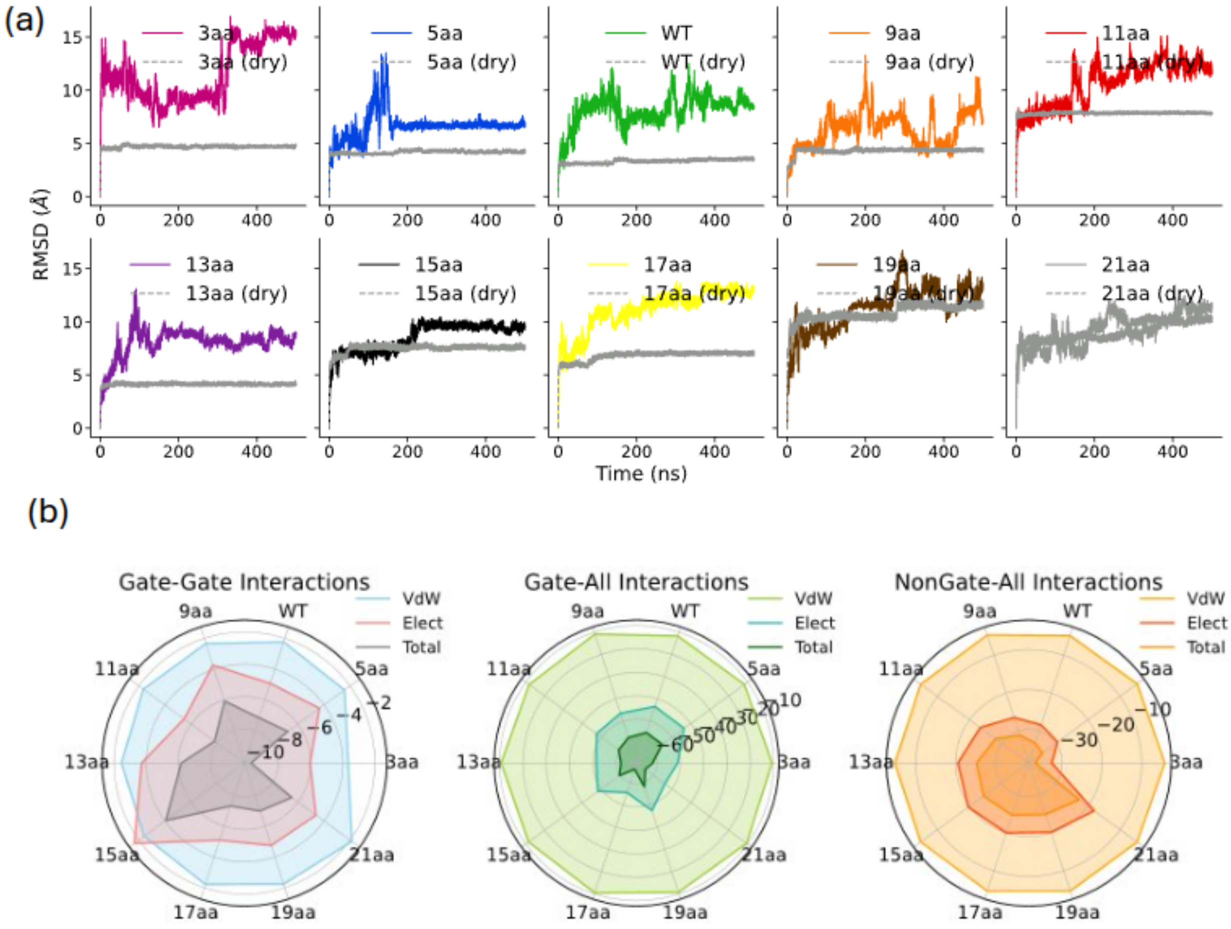

**Supplementary Figure S4**: (a)–(c) Snapshots of conformations at different MD frames show structural changes for 15aa in aqueous medium. (d)–(f) show conformations of 19aa at different time frames.

.

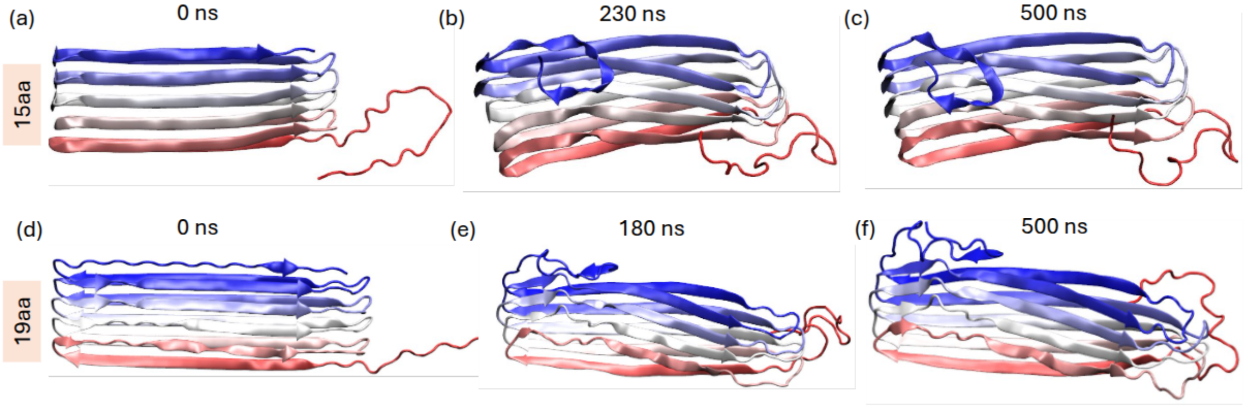

**Supplementary Figure S5:** (a) Radial distribution function (RDF) of polar (including charged) and nonpolar residues of core β-strands with oxygen atoms in water. (b) Normalized density of water as a function of distance from the polar and nonpolar residues of the protein.

**Hydration of the polar and nonpolar residues:** Radial distribution functions (RDFs) were calculated for polar (including charged) and nonpolar residues of core β-strands with respect to oxygen atoms in water for all variants, as shown in **Fig. S5a**. The RDF peaks indicate that polar residues are generally more hydrated than nonpolar residues across all variants, with the exception of the 5aa variant, where nonpolar residues exhibit hydration levels comparable to those of polar residues. To further investigate, we computed the normalized water density around polar and nonpolar residues as a function of distance in **Fig. S5b**. The first hydration shell, comprising peaks at 2.5 Å and 2.8 Å, is predominantly contributed by polar residues, while the second hydration peak at approximately 6.0 Å is equally influenced by both polar and nonpolar residues.

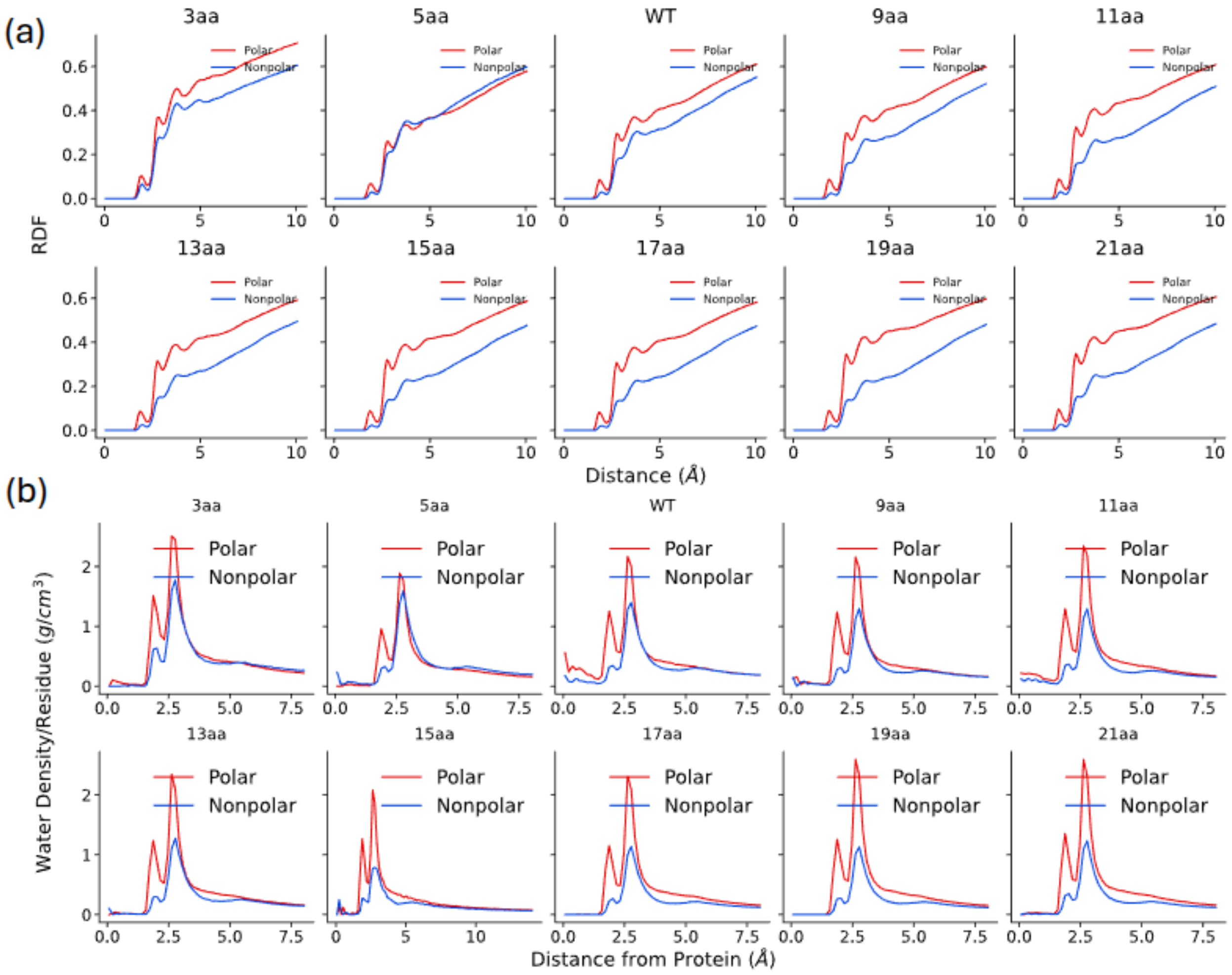

**Supplementary Figure S6:** Average persistence length of individual β-strands (β1 to β10) as a function of simulation time. The standard deviations are shown in braces. The average of core β-strands (β2 to β9) is shown in the final column.

**Persistence length of individual β-strands:** The persistence length of individual β-strands within a β-solenoid protein was calculated as a block average over 25 ns time windows, evaluated as a function of time (see **Fig. S6**). A positive correlation was observed between the persistence length and the length of the β-strands, suggesting that longer strands exhibit greater resistance to bending. Among the β-strands analyzed, the C-terminal β-10 strand consistently demonstrated the lowest persistence length, with values decaying over time. This observation aligns with its elevated root-mean-square fluctuation (RMSF) values, indicative of higher flexibility. In contrast, β-strands located in the central region of the β-solenoid exhibited greater persistence lengths, which can be attributed to an increased probability of hydrogen bond formation with adjacent strands.

| | Individual β-strand Persistence Length | | | | | | | | | | |
|---|---|---|---|---|---|---|---|---|---|---|---|
| Species | β1 | β2 | β3 | β4 | β5 | β6 | β7 | β8 | β9 | β10 | Avg (β2:β9) |
| 3aa | 5.14 (0.78) | 4.24 (2.79) | 9.96 (4.15) | 4.85 (0.66) | 5.51 (0.73) | 4.78 (0.65) | 5.83 (0.81) | 18.04 (6.33) | 5.71 (1.41) | 6.11 (2.28) | 7.367 |
| 5aa | 14.07 (1.66) | 16.25 (0.81) | 13.72 (0.30) | 40.88 (0.95) | 9.20 (0.14) | 43.58 (2.18) | 13.66 (0.76) | 16.30 (1.10) | 18.63 (1.34) | 15.12 (3.88) | 21.528 |
| WT | 21.47 (1.35) | 41.81 (2.55) | 21.11 (0.58) | 28.95 (1.50) | 65.63 (1.11) | 75.60 (3.00) | 76.09 (3.85) | 39.26 (1.65) | 56.55 (7.99) | 10.79 (14.65) | 50.623 |
| 9aa | 68.88 (5.59) | 41.43 (1.79) | 84.26 (1.10) | 36.78 (0.84) | 41.30 (0.69) | 46.12 (8.00) | 89.33 (2.66) | 76.83 (7.06) | 41.21 (6.45) | 24.61 (16.66) | 57.157 |
| 11aa | 37.75 (21.00) | 94.90 (10.14) | 35.24 (1.29) | 53.14 (3.84) | 55.12 (1.58) | 38.58 (2.97) | 104.54 (5.60) | 56.90 (19.92) | 88.44 (9.16) | 10.74 (13.13) | 65.857 |
| 13aa | 50.37 (29.00) | 85.37 (13.88) | 111.18 (8.39) | 60.73 (2.17) | 91.39 (3.65) | 87.88 (8.30) | 97.83 (3.96) | 89.58 (6.87) | 101.05 (3.24) | 7.17 (11.28) | 90.628 |
| 15aa | 99.51 (9.04) | 89.77 (13.97) | 109.61 (3.57) | 75.82 (4.26) | 116.94 (2.79) | 96.58 (7.80) | 57.12 (1.99) | 85.24 (8.74) | 113.36 (7.81) | 31.15 (33.20) | 93.054 |
| 17aa | 124.33 (10.24) | 85.19 (16.92) | 147.57 (3.33) | 100.93 (17.18) | 142.09 (3.37) | 109.10 (9.52) | 143.85 (4.23) | 121.00 (8.27) | 138.25 (7.77) | 11.89 (16.39) | 123.498 |
| 19aa | 143.93 (10.77) | 85.27 (7.44) | 147.01 (3.31) | 93.15 (14.33) | 139.81 (3.79) | 105.63 (1.88) | 135.72 (4.19) | 118.81 (11.78) | 134.51 (9.64) | 19.60 (37.21) | 119.990 |
| 21aa | 156.56 (18.96) | 84.53 (12.02) | 122.84 (10.42) | 82.75 (8.57) | 155.94 (3.19) | 83.96 (11.93) | 142.60 (7.63) | 102.53 (17.73) | 132.07 (16.21) | 54.81 (36.73) | 113.403 |

**Supplementary Figure S7:** Snapshots of the conformations (front and back) of all the variants. The positively and negatively charged residues are highlighted in red and green, respectively.

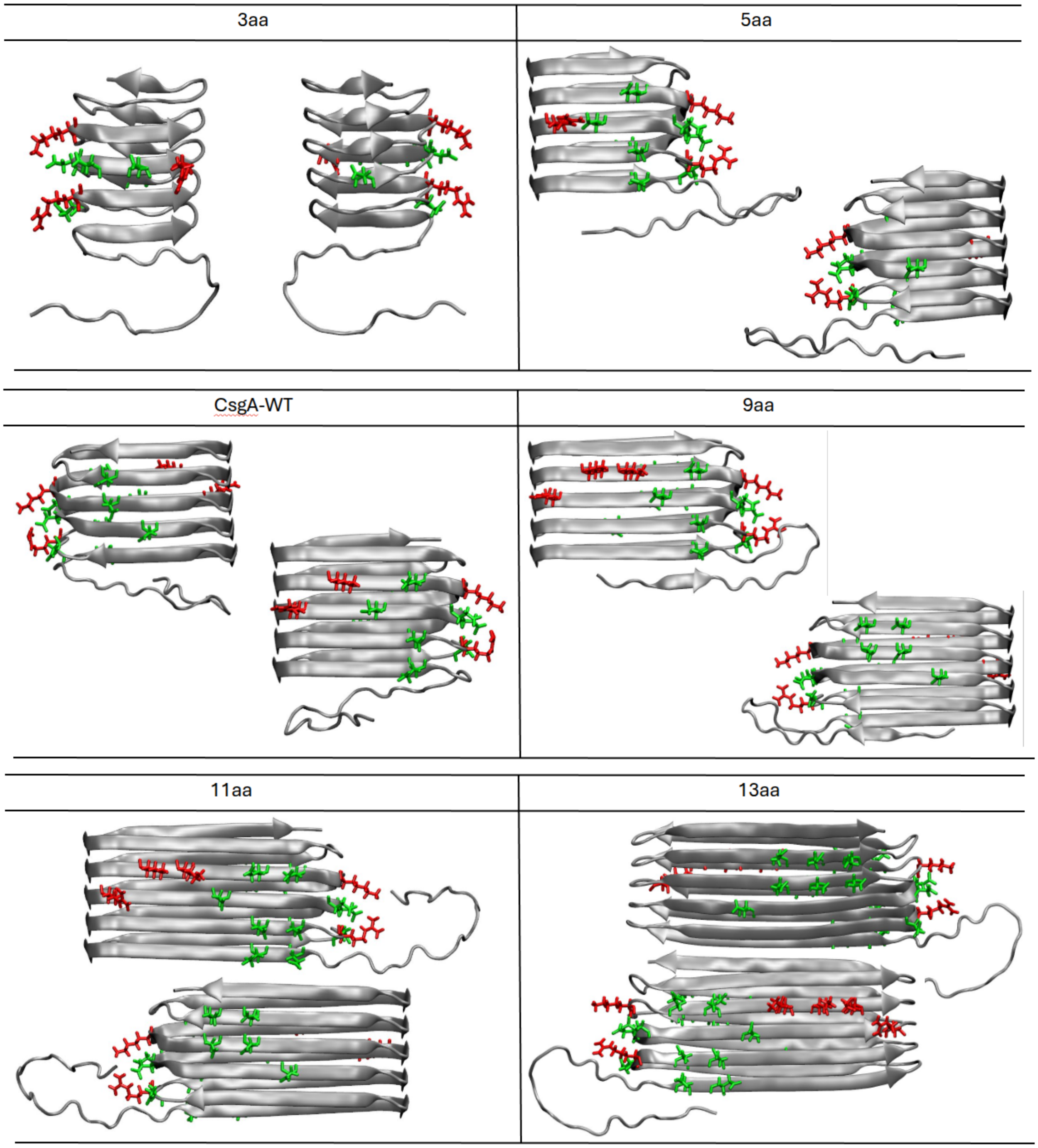

| 15aa | 17aa |
|---|---|
| 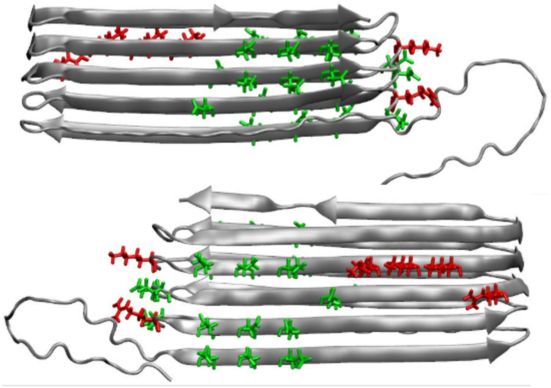 | 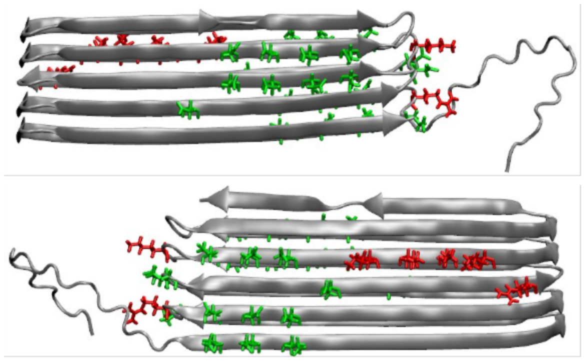 |
| **19aa** | **21aa** |
| 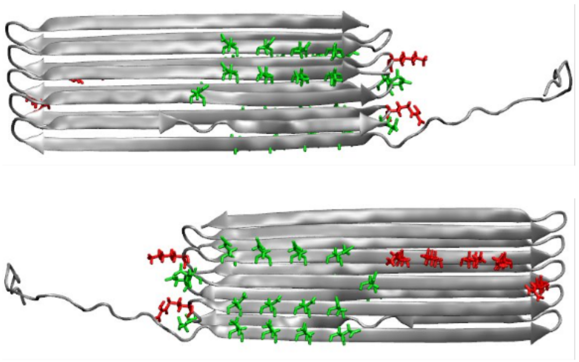 | 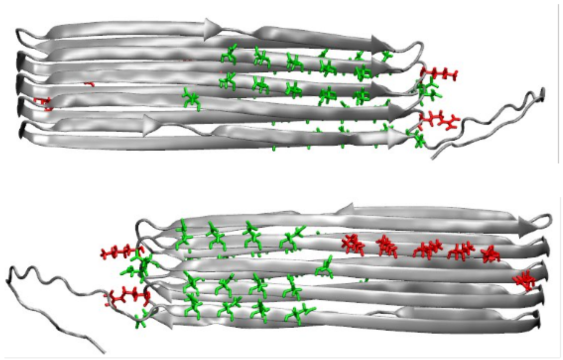 |

**Supplementary Figure S8:** The van der Waals, electrostatics, and total interaction energies of (a) polar to polar (including the charged residues), (b) charged to charged, (c) non-polar to non-polar, and (d) polar to non-polar residues. The inset of (a) shows the interaction energies among charged residues only.

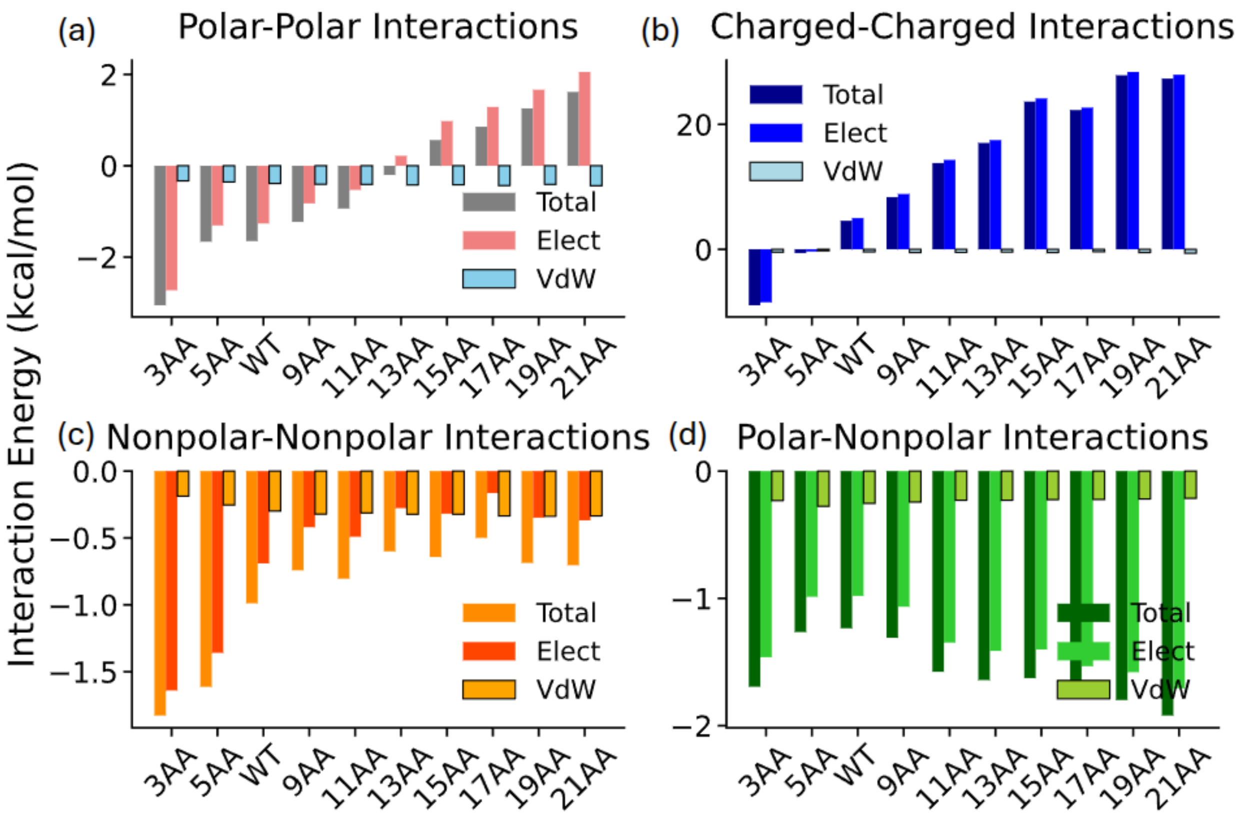

**Supplementary Figure S9:** The van der Waals, electrostatics, and total interaction energies for the first 50 ns for (a) gate-to-gate, (b) gate-to-all, (c) nongate-to-all, (d) gate-to-water, and (e) nongate-to-water.

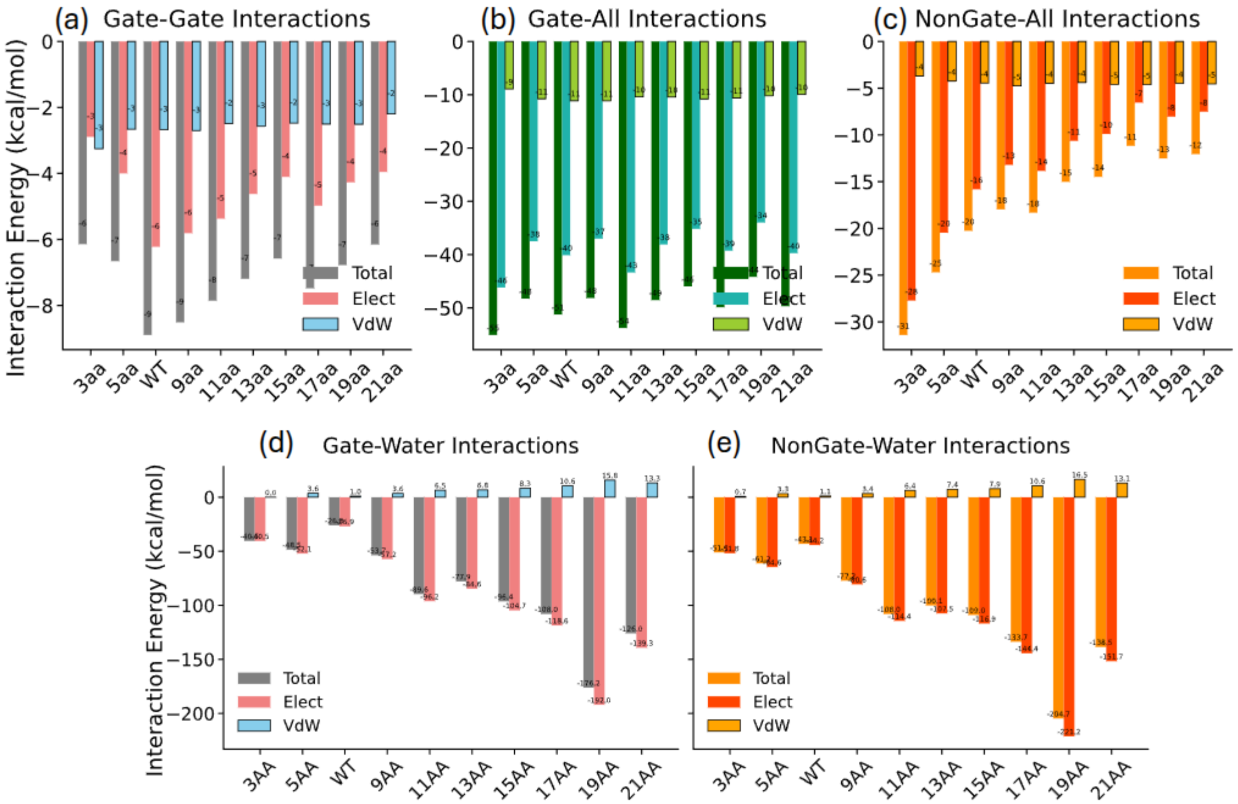

**Supplementary Figure S10:** Radial distribution function (RDF) between protein backbone heavy atoms and oxygen atoms of water as a function of radial distance for all the CsgA variants in the first 50 ns of MD simulation.

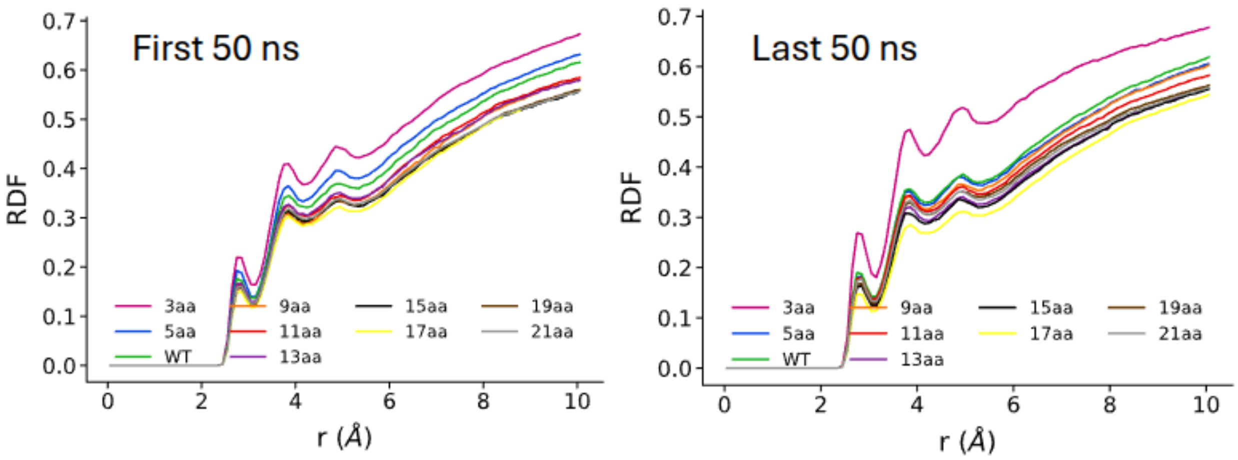

**Supplementary Figure 11:** Tetrahedral orientational water order parameter for all the CsgA variants in the first 50 ns of MD simulation. The bulk water order is shown in the dotted grey line for comparison.

**Tetrahedral orientational order of water:** To characterize the extent of water ordering in the vicinity of the protein, The normalized tetrahedral orientational order parameter (q) for water was calculated over the first 50 ns (see **Fig. S11**) and the final 50 ns (see **Fig. S4b** in the main manuscript) of the MD simulation. This parameter, which quantifies the tetrahedral arrangement of water molecules, was computed using the four nearest oxygen neighbors of each water molecule. The data were smoothed using the Savitzky-Golay filter [18] with a third-degree polynomial and a window length of 31 to precisely determine peak positions. The value of q ranges from 0 (ideal gas) to 1 (perfect tetrahedron). Our results show a water ordering parameter of approximately q ≈ 0.4 near the protein surface, while the bulk TIP3P water exhibits an order parameter of approximately 0.5, consistent with previously reported literature values [19]. Near the amyloids, the OTO peak values were found to shift leftward for the last 50 ns of simulation, ranging from 0.34 to 0.39, indicating a distinct water ordering compared to bulk water. Specifically, the 5aa and 15aa variants exhibited lower ordering, with peaks at 0.3, whereas the peaks for the other variants remained close to 0.38 (see Supplementary **Table S2**). The order parameters for both the first and last 50 ns are summarized in **Table S11**.

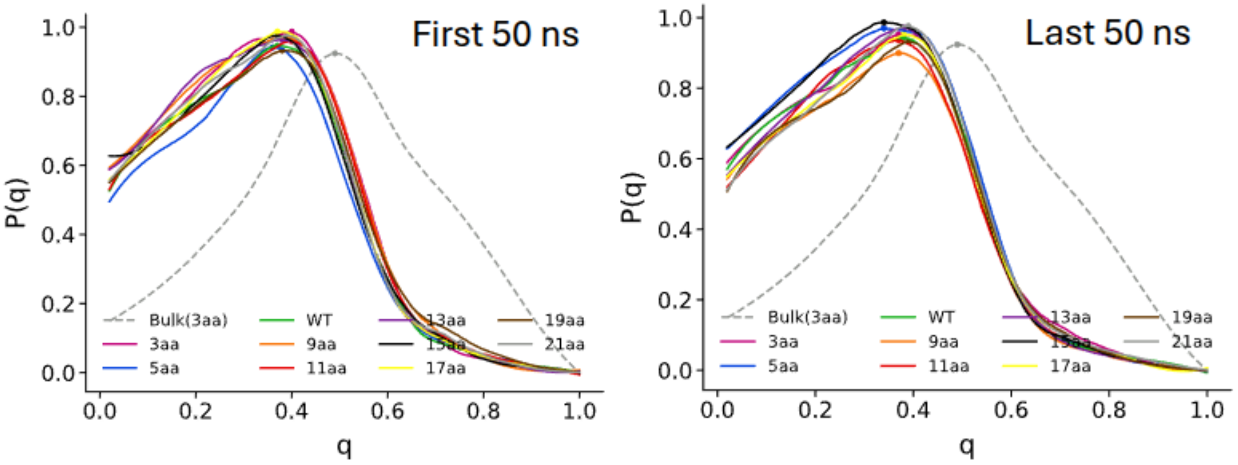

**Supplementary Table S11:** The table shows the tetrahedral orientational water order parameter (q) and the normalized distribution peak height P(q) for the first 50 ns and the last 50 ns of the simulation.

| First 50 ns | | |
|---|---|---|
| **Variants** | **q** | **P(q)** |
| 3aa | 0.4 | 0.987 |
| 5aa | 0.36 | 0.941 |
| CsgA_WT_7AA | 0.37 | 0.947 |
| 9aa | 0.39 | 0.969 |
| 11aa | 0.4 | 0.960 |
| 13aa | 0.38 | 0.964 |
| 15aa | 0.37 | 0.976 |
| 17aa | 0.37 | 0.987 |
| 19aa | 0.38 | 0.933 |
| 21aa | 0.39 | 0.975 |
| Bulk | 0.49 | 0.924 |

| Last 50 ns | | |
|---|---|---|
| **Variants** | **q** | **P(q)** |
| 3aa | 0.37 | 0.953 |
| 5aa | 0.34 | 0.970 |
| CsgA_WT_7AA | 0.38 | 0.944 |
| 9aa | 0.37 | 0.899 |
| 11aa | 0.37 | 0.936 |
| 13aa | 0.38 | 0.971 |
| 15aa | 0.34 | 0.986 |
| 17aa | 0.38 | 0.952 |
| 19aa | 0.39 | 0.932 |
| 21aa | 0.39 | 0.976 |
| Bulk | 0.49 | 0.924 |

**Supplementary Figure S12:** The comparison of RMSD of the C-alpha atoms of the core β-solenoids (in color) with the RMSD in the absence of partial charges in charged residues (Arg, Lys, Asp, Glu) in cyan and for all residues in grey.

**Role of charged residues in β-solenoid stability:** To investigate the role of charged residues in the stability of β-solenoid proteins, we performed molecular dynamics simulations in which partial charges were artificially neutralized for (a) only the charged residues and (b) all residues. In both scenarios, the systems were charge-neutral, and no counter-ions were included. The root-mean-square deviation (RMSD) of the β-solenoid structures was monitored over 100 ns and compared to simulations with intact charges. **Fig. S3a** illustrates the case where only charged residues were neutralized (shown in grey). The results indicate that, except for the 3aa and 9aa variants, the RMSD was lower compared to the charged systems, suggesting that repulsion between like-charged residues may slightly destabilize the protein structure. This effect was particularly pronounced in the 5aa, 11aa, and 13aa variants, where neutralizing charged residues markedly enhanced stability. In contrast, **Fig. S3b** shows the extreme case where all partial charges were neutralized, resulting in a ubiquitous increase in RMSD and notable loss of secondary structure across all variants.

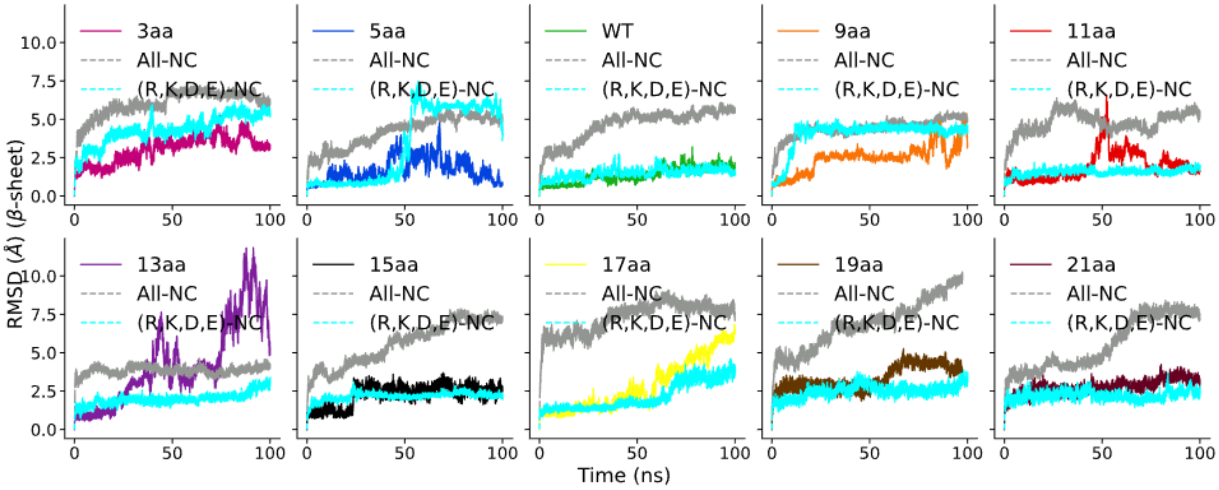

**Supplementary Figure S13:** The comparison of β-sheet percentage as a function of MD time for all the CsgA variants and CsgA WT (in green). The data is smoothed using the Savitzky-Golay filter with a second-degree polynomial and a window length of 51.

**Percentage of β-strands in time:** The normalized percentage of β-sheet content was calculated across the entire 500 ns molecular dynamics simulation trajectory shown in **Fig. S13**. The β-sheet percentage was normalized relative to the initial frame, defined as 100%. For all variants except 3aa, the β-sheet content remained stable at approximately 80% throughout the simulation duration. In contrast, the 3aa variant exhibited a substantial reduction in β-sheet content, confirming the instability of its secondary structure beyond 50 ns.

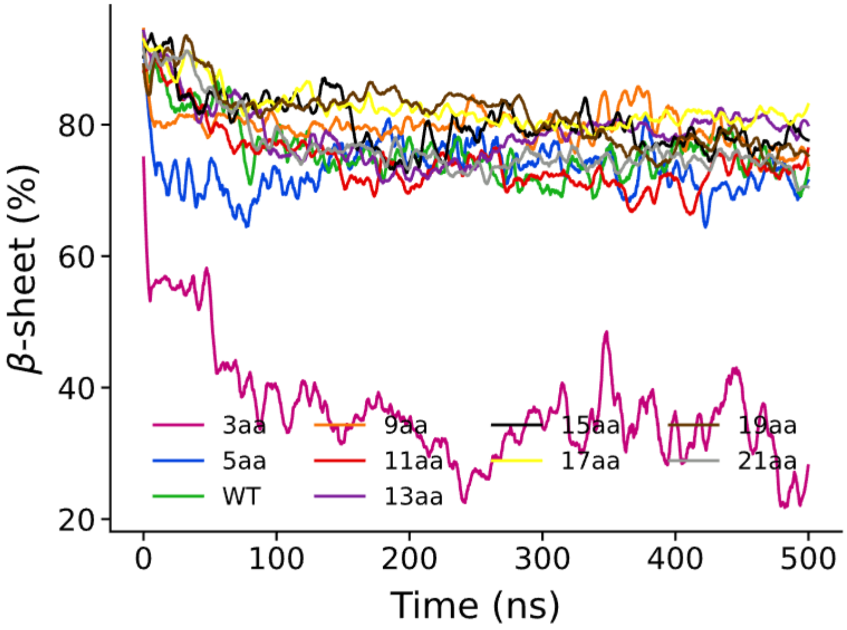

**Supplementary Figure S14:** Comparison of residence time of ions near 6 Å of the charged residues. The y-axis represents normalized residence time, normalized with respect to the average residence time of ion near neutral residues. The green, red, and blue colors are chosen to highlight negatively (Asp and Glu), positively (Arg and Lys), and neutrally charged residues.

**Ion residence time:** The residence times of ions within 6 Å of protein residues were calculated over the final 50 ns of the molecular dynamics (MD) trajectory. Residence times were normalized by the average residence time of ions near neutral residues, enabling a comparative analysis of ion retention near charged residues relative to neutral ones, shown in **Fig. S14**. Given the net negative charge of all variants (see Table S1), only counterbalancing sodium ions were present in the simulation. The results demonstrate that sodium ions exhibit significantly longer residence times near negatively charged residues (green) compared to positively charged (red) or neutral residues (blue). For larger variants (13aa to 21aa), which possess a larger number of charged residues, ion residence times increased significantly near negatively charged patches (see **Fig. S7**). This behavior may mitigate inter-charge repulsion among closely packed negatively charged residues, potentially stabilizing the secondary structure in these larger variants, and may serve as a sodium-binding pocket like GPCRs [1].

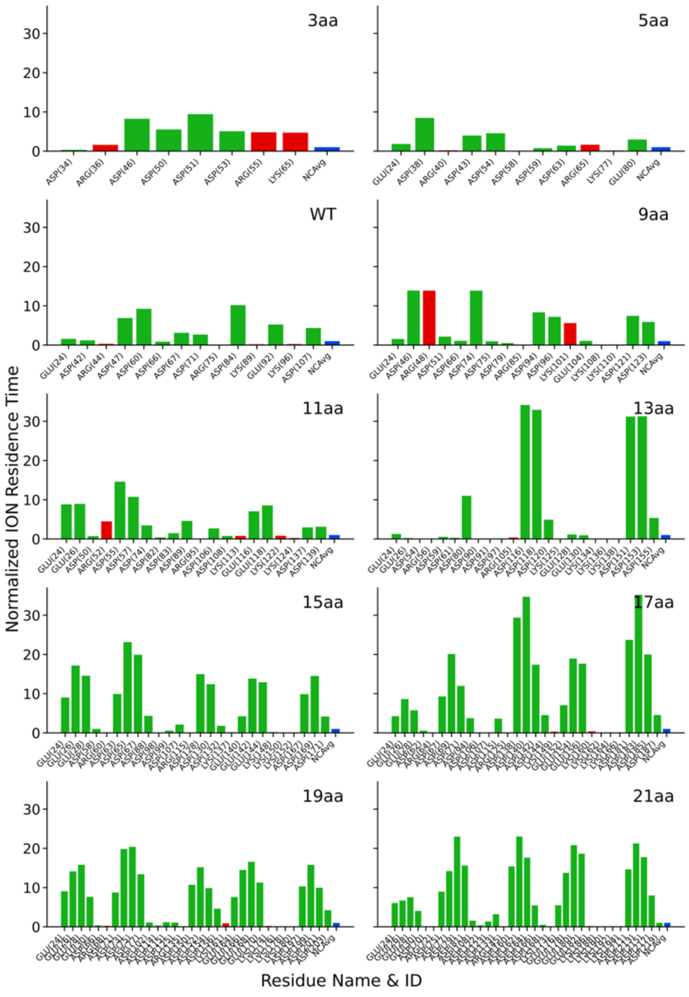

**Supplementary Table S3:** The orientational tetrahedral order parameter (OTO) for CsgA WT and all the variants.

| Table for per Residue Interaction Energies (within 12 A) | | | | | | | | | |
|---|---|---|---|---|---|---|---|---|---|
| Species | Gate-Gate (CR-CR) | | | Gate-All (CR-All) | | | NonGate-All (NonCR-All) | | |
| | VdW | Elect | Total | VdW | Elect | Total | VdW | Elect | Total |
| 3aa | -1.599 | -1.916 | -3.515 | -6.700 | -46.077 | -52.777 | -3.614 | -27.730 | -31.344 |
| 5aa | -2.629 | -4.321 | -6.949 | -10.715 | -38.206 | -48.920 | -4.354 | -23.860 | -28.214 |
| WT | -2.330 | -4.829 | -7.158 | -10.314 | -39.506 | -49.820 | -4.504 | -17.114 | -21.617 |
| 9aa | -2.409 | -5.067 | -7.476 | -10.391 | -38.642 | -49.034 | -4.533 | -14.310 | -18.842 |
| 11aa | -2.044 | -4.365 | -6.409 | -9.568 | -42.711 | -52.279 | -4.441 | -16.975 | -21.416 |
| 13aa | -2.574 | -4.005 | -6.579 | -10.522 | -35.797 | -46.319 | -4.575 | -13.544 | -18.119 |
| 15aa | -2.125 | -3.985 | -6.110 | -10.235 | -40.188 | -50.423 | -4.645 | -11.609 | -16.254 |
| 17aa | -2.440 | -3.993 | -6.432 | -11.157 | -36.192 | -47.350 | -4.688 | -9.544 | -14.232 |
| 19aa | -1.694 | -3.524 | -5.218 | -9.517 | -42.297 | -51.814 | -4.454 | -10.708 | -15.162 |
| 21aa | -1.785 | -3.589 | -5.374 | -9.086 | -43.779 | -52.865 | -4.498 | -10.008 | -14.506 |

**Supplementary Table S4:** Normalized residue-residue van der Waals (VdW), electrostatic, and total interaction energies for gate-gate, gate-all, and non-gate-to-all interactions for the CsgA variants.

| Table for per Residue Interaction Energies (within 12 A) - DRY RUN | | | | | | | | | |
|---|---|---|---|---|---|---|---|---|---|
| Species | Gate-Gate (CR-CR) | | | Gate-All (CR-All) | | | NonGate-All (NonCR-All) | | |
| | VdW | Elect | Total | VdW | Elect | Total | VdW | Elect | Total |
| 3aa | -3.667 | -6.042 | -9.709 | -10.721 | -51.222 | -61.943 | -4.176 | -32.648 | -36.824 |
| 5aa | -2.430 | -4.352 | -6.782 | -11.675 | -43.635 | -55.309 | -4.685 | -29.483 | -34.168 |
| WT | -2.240 | -4.933 | -7.173 | -11.710 | -43.453 | -55.163 | -4.849 | -28.393 | -33.242 |
| 9aa | -2.340 | -3.748 | -6.088 | -10.909 | -46.481 | -57.390 | -4.664 | -26.485 | -31.149 |
| 11aa | -2.341 | -5.472 | -7.813 | -11.776 | -47.535 | -59.311 | -4.809 | -23.324 | -28.133 |
| 13aa | -2.468 | -3.719 | -6.187 | -10.649 | -51.458 | -62.106 | -4.706 | -20.587 | -25.293 |
| 15aa | -2.407 | -1.660 | -4.067 | -11.447 | -48.263 | -59.710 | -4.762 | -19.606 | -24.368 |
| 17aa | -2.223 | -5.092 | -7.315 | -10.497 | -55.539 | -66.036 | -4.669 | -20.034 | -24.703 |
| 19aa | -2.240 | -4.750 | -6.991 | -10.756 | -47.444 | -58.200 | -4.667 | -20.220 | -24.887 |
| 21aa | -1.864 | -4.623 | -6.487 | -10.439 | -53.826 | -64.266 | -4.709 | -18.063 | -22.772 |

**Supplementary Table S5:** Normalized residue-residue van der Waals (VdW), electrostatic, and total interaction energies for gate-gate, gate-all, and non-gate-to-all interactions for the CsgA variants in the absence of water.

| Table for Residue Water Interaction Energies | | | | | | |
|---|---|---|---|---|---|---|
| Species | Gate-Water (CR-Water) | | | NonGate-Water (NonCR-Water) | | |
| | VdW | Elect | Total | VdW | Elect | Total |
| 3aa | 0.034 | -40.488 | -40.454 | 0.736 | -51.809 | -51.073 |
| 5aa | 3.638 | -52.142 | -48.504 | 3.341 | -64.581 | -61.240 |
| WT | 0.958 | -26.941 | -25.983 | 1.087 | -44.201 | -43.114 |
| 9aa | 3.583 | -57.248 | -53.665 | 3.430 | -80.630 | -77.200 |
| 11aa | 6.545 | -96.194 | -89.649 | 6.374 | -114.422 | -108.049 |
| 13aa | 6.772 | -84.644 | -77.872 | 7.375 | -107.496 | -100.121 |
| 15aa | 8.315 | -104.673 | -96.358 | 7.903 | -116.866 | -108.963 |
| 17aa | 10.607 | -118.648 | -108.041 | 10.635 | -144.353 | -133.717 |
| 19aa | 15.829 | -192.032 | -176.203 | 16.498 | -221.211 | -204.713 |
| 21aa | 13.308 | -139.269 | -125.961 | 13.137 | -151.684 | -138.547 |

**Supplementary Table S6:** Normalized residue-water van der Waals (VdW), electrostatic, and total interaction energies for gate-water and non-gate-to-water interactions for the CsgA variants.

| Table for per Residue Interaction Energies (within 12 A) - Polar/Non-Polar | | | | | | | | | | | | | |
|---|---|---|---|---|---|---|---|---|---|---|---|---|---|
| Species | Polar-Water | | | NonPolar-Water | | | Charged-Water | | | Polar | NonPolar | Charged | Total |
| | VdW | Elect | Total | VdW | Elect | Total | VdW | Elect | Total | Count | Count | Count | Count |
| 3aa | 0.125 | -45.232 | -45.107 | 0.815 | -37.777 | -36.962 | 2.369 | -116.240 | -113.871 | 45 | 38 | 8 | 91 |
| 5aa | 3.308 | -59.705 | -56.397 | 3.674 | -57.863 | -54.189 | 4.746 | -108.623 | -103.877 | 50 | 50 | 11 | 111 |
| WT | 0.481 | -33.724 | -33.243 | 0.468 | -29.031 | -28.563 | 4.167 | -126.292 | -122.125 | 55 | 62 | 14 | 131 |
| 9aa | 2.563 | -62.236 | -59.673 | 3.266 | -61.123 | -57.857 | 7.653 | -206.091 | -198.438 | 60 | 74 | 17 | 151 |
| 11aa | 5.143 | -96.308 | -91.165 | 5.481 | -83.695 | -78.214 | 11.555 | -262.732 | -251.177 | 65 | 86 | 20 | 171 |
| 13aa | 5.973 | -85.547 | -79.574 | 7.316 | -91.437 | -84.121 | 11.116 | -200.858 | -189.742 | 70 | 98 | 23 | 191 |
| 15aa | 7.223 | -101.471 | -94.247 | 7.007 | -89.859 | -82.851 | 12.237 | -235.554 | -223.317 | 75 | 110 | 26 | 211 |
| 17aa | 9.615 | -122.456 | -112.841 | 10.397 | -122.070 | -111.673 | 13.884 | -264.213 | -250.329 | 80 | 122 | 29 | 231 |
| 19aa | 15.581 | -196.445 | -180.864 | 16.387 | -191.107 | -174.720 | 20.081 | -382.825 | -362.744 | 85 | 134 | 32 | 251 |
| 21aa | 12.449 | -136.927 | -124.477 | 12.876 | -138.001 | -125.125 | 15.395 | -226.438 | -211.044 | 90 | 146 | 35 | 271 |

**Table S7:** Normalized residue-water van der Waals (VdW), electrostatic, and total interaction energies for polar residue-to-water (excluding the charged residues), nonpolar residue-to-water, and charged residue-to-water interactions for the CsgA variants.
